# A new resource of clonal pluripotent human stem cell lines exhibiting inter- and intra-embryo consistency and variability

**DOI:** 10.1101/2025.06.02.657331

**Authors:** Stanley E. Strawbridge, Lawrence E. Bates, Connor Ross, Kenneth A. Jones, Takuya Azami, Tim Lohoff, Maike Paramor, Vicki Murray, Ana Luíza Cidral, James Clarke, Maria Rostovskaya, Ge Guo, Jennifer Nichols

## Abstract

Human naive pluripotent stem cells can generate all somatic tissues and extra-embryonic components of the blastocyst. We derived multiple clonal naive pluripotent stem cell lines from individual embryos by physical separation of inner cell mass cells and subsequent individual expansion of each resulting dome-shaped colony, providing the foundation for a resource to investigate intra- and inter-embryo variation. Twenty lines were derived from ten embryos donated from nine couples. While differences between lines are observed, the overarching pluripotency circuitry is preserved in each. They can differentiate into extra-embryonic lineages and readily acquire post-implantation pluripotent identity when exposed to culture conditions driving *in vitro* capacitation, subsequently to generate derivatives of the three germ layers: ectoderm, mesoderm and endoderm. Some lines exhibit intra-chromosomal amplification and deletions and are therefore anticipated to provide a valuable, accessible system for modelling chromosomal mosaicism and its potential consequences using chimeric organoids, such as blastoids and gastruloids.

## INTRODUCTION

Within 4-6 days of fertilisation the human embryo undergoes cavitation to form a blastocyst comprising an outer layer of trophectoderm, precursor of the placenta marked by expression of GATA3 (Figure 1A), surrounding a bipotent progenitor cluster of cells known as the ‘inner cell mass’ (ICM), expressing OCT4 and PDGFRa (Corujo-Simon et al., 2023). The ICM then gradually specifies into NANOG+ epiblast, founder of the fetus and source of pluripotent stem cells, and GATA4+ hypoblast that will produce anteriorising signals, and subsequently form the yolk sac. By the mid-late blastocyst stage, at around D6, the ICM will have fully segregated into naïve pluripotent epiblast and hypoblast, at which point epiblast cells acquire the potential to be captured *in vitro* as naïve pluripotent stem cell lines (nPSCs) (Guo et al., 2016). Chromosomal mosaicism is frequently observed in human embryos, resulting from emergence of aneuploidy during early development (Evsikov and Verlinsky, 1998; Johnson et al., 2010). Aneuploidies can arise at cleavage stages (Babariya et al., 2017), with effects persisting until after birth, potentially resulting in genetic disease states. While it has been proposed that some aneuploidy-exhibiting cells are eliminated from the ICM during preimplantation development (Magli et al., 2000), a mechanism remains to be defined. The potential to separate and expand euploid and aneuploid cells from the same genetic background in culture may lead to better understanding of the consequences when specific aneuploid cells persist during development.

**Figure 1.**
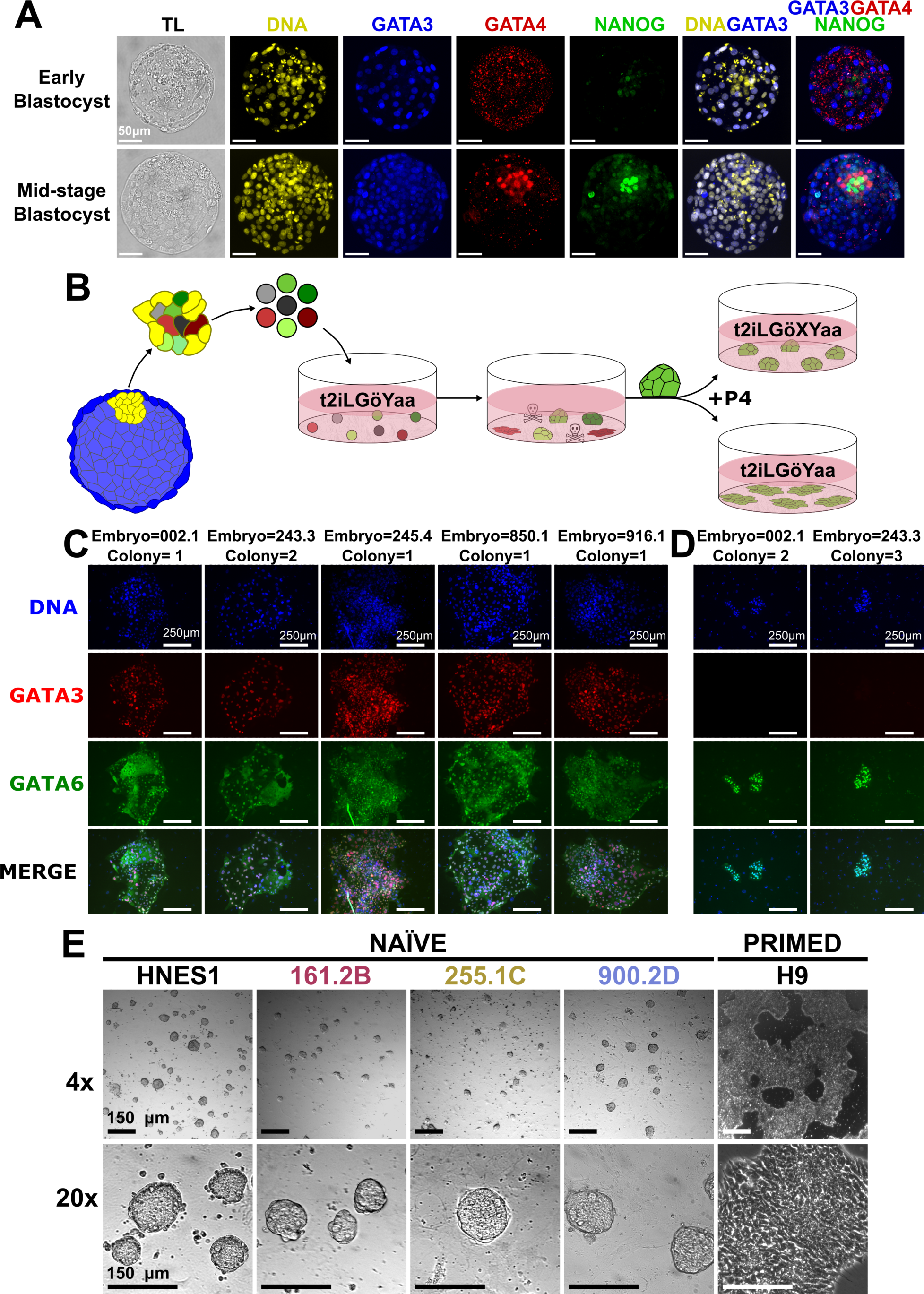
Establishing clonal human naïve pluripotent stem cell lines. (A) Immunofluorescence of early and mid-stage human blastocysts stained for DNA (DAPI; yellow) and lineage markers indicating trophoblast (GATA3; blue), hypoblast (GATA4; red), and epiblast (NANOG; green). (B) Schematic of clonal naïve pluripotent stem cell (nPSC) line derivation. The inner cell mass (yellow) is isolated from trophectoderm (blue) by immunosurgery, disaggregated into single cells or small clumps and cultured in t2iLGöYaa. Dome-shaped colonies (green) are disaggregated and passaged in t2iLGöYaa until passage 4, when the media is supplemented with XAV939 to maintain domed morphology and naïve pluripotency. (C) Immunofluorescence of spontaneous GATA3+ (red)/ GATA6+ (green) trophoblast-like differentiation; (D) GATA3-/GATA6+ hypoblast-like differentiation during early passage expansion (prior to addition of Xav939). The Embryo nomenclature is donor-embryo, where 002.1 indicates the anonymised donor code “002” and embryo “1” from donor 002. (E) Bright field images of previously derived HNES1 and our case study clonal lines (161.2B, 255.1C, and 900.2D) in nPSC culture conditions, and the previously derived conventional primed PSC line H9.

The first pluripotent embryonic stem cell lines, defined by their capacity to self-renew indefinitely and retain the potential to differentiate into any cell in the body, were derived from mouse embryos using medium containing serum and a ‘feeder’ layer of mitotically inactivated fibroblasts (Evans and Kaufman, 1981; Martin, 1981). These cells, subsequently termed ‘naïve pluripotent’, exist in appropriate culture conditions as dome-shaped colonies with indistinct boundaries between cells. Applying a similar approach, but with the addition of fibroblast growth factor (FGF), derivation of human pluripotent stem cell lines from donated embryos generated by *in vitro* fertilisation was achieved (Thomson et al., 1998). Interestingly, although originating from blastocysts, these lines more closely resembled self-renewing populations, known as ‘epiblast stem cells’ (EpiSCs) derived from isolated epiblasts of postimplantation mouse embryos (Brons et al., 2007; Tesar et al., 2007), which exhibit an epithelial morphology and more restricted differentiation potential, and therefore referred to as ‘primed pluripotent’ (Nichols and Smith, 2009). It is postulated that during the derivation process, explanted human blastocysts are relatively unconstrained from transitioning into the epithelialized state characteristic of the post implantation epiblast compared with mouse embryos.

Experimental blockade of FGF signalling via MEK/ERK inhibition applied to mouse embryos from the morula stage prevents formation of hypoblast (known as ‘primitive endoderm’ in mouse) (Nichols et al., 2009), whereas more potent inhibition of FGF signalling is required to impede formation of human hypoblast (Dattani et al., 2024). To minimise transfer of inductive FGF signals from the emerging hypoblast to the epiblast, and avoid the potentially detrimental effects of added inhibitors, in this study we physically separated disaggregated ICM cells to facilitate derivation of clonal naïve pluripotent stem cells (nPSCs) from human embryos. We developed this system previously by isolating ICMs using immunosurgery (Solter and Knowles, 1975) from D6 human blastocysts (Guo et al., 2016). In that study, colonies from disaggregated ICMs with domed morphology were pooled for each individual embryo to maximise the likelihood of successful expansion of lines retaining naïve pluripotent status. We subsequently enhanced the protocol to enable derivation of several lines from the same embryo by expanding each dome-shaped colony individually. A detailed protocol for this method has been published (Strawbridge et al., 2022).

For the present study we undertook lineage differentiation analysis of a subset of the new lines, including capacitation and directed differentiation to the three germ layers and extra-embryonic tissues. Karyotypic characterisation revealed some intra-embryo variations. In total we derived and successfully expanded twenty clonal nPSC lines from ten embryos donated by nine couples, which we present as a resource to be available for the research community.

## RESULTS

### Physical separation of inner cell mass cells enables derivation of multiple naïve pluripotent stem cell clones per embryo

Naïve pluripotent stem cell line derivations were conducted using a similar protocol to the one we devised previously by isolating ICMs using immunosurgery (Solter and Knowles, 1975) and disaggregating them into single cells or small clusters (Guo et al., 2016) (Figure 1B). We instigated two major improvements. Firstly, colonies with the dome-shape morphology characteristic of the naïve pluripotent state were propagated and expanded individually to generate clonal lines (Strawbridge et al., 2022). The second modification was addition of the Tankyrase inhibitor, XAV939, to the original t2iLGöYaa medium at passage 4 in order to supress differentiation (Dattani et al., 2022). Spontaneous colony epithelialisation of nascent cell lines and induction of markers of trophoblast (Figure 1C) or hypoblast (Figure 1D) could be inhibited by addition of XAV939 to t2iLGöYaa medium. In the absence of XAV939, nPSCs can readily differentiate into trophoblast (Guo et al., 2016; Io et al., 2021), hypoblast (Dattani et al., 2024), extra-embryonic endoderm (Linneberg-Agerholm et al., 2019), and the entire blastocyst structure in the form of ‘blastoids’ (Kagawa et al., 2022; Liu et al., 2021; Yanagida et al., 2021).

Derivations were attempted with 84 viable embryos, yielding 20 lines from 10 embryos from 9 genetically distinct donors (Table S1). This includes one set of 4 lines, 2 sets of 3 lines, 3 sets of 2 lines, and 4 single lines from individual embryos. We used nomenclature of donor-embryo-clone identifiers, for example, for the line labelled 161.2B, “161” is the anonymised donor code, “2” indicates the second embryo from donor 161, and “B” indicates the second clone expanded from the initial plating of the ICM cells from embryo 2. Three lines: 161.2B, 255.1C, and 900.2D (Table S1) were randomly selected from different genetic backgrounds (Figure 1E) as a case study for subsequent basic differentiation analysis.

### Hallmarks of naïve pluripotency are conserved between genetic backgrounds of clonal nPSCs

We explored the potential of our three case study lines to differentiate by driving developmental progression to the primed state via *in vitro* capacitation (Rostovskaya et al., 2019) (Figure 2A) and compared morphology and expression of key markers at both RNA and protein levels. The expected morphological change from discrete, domed naïve colonies (Figure 1E) to epithelial monolayer primed colonies was observed (Figure 2B). While naïve markers KLF4 and KLF17 were down-regulated, pan-pluripotency markers OCT4 (*POU5F1*) and NANOG were maintained during the transition (Figure 2C to J). RNA levels of *OCT4* and *SOX2* (Figure 2D and K) remained constant between the two states, as expected (p-value = 0.5 and 0.8, respectively; one-tailed Wilcoxon Rank Sum, n=3), while NANOG levels were lower in the primed state (p-value = 0.05, one-tailed Wilcoxon Rank Sum, n=3). Expression of naïve markers *KLF4*, *KLF17*, *GDF3*, *NODAL*, *SUSD2*, *TFAP2C*, and *TFCP2L1* were all higher in the naïve than the primed state, as assessed using RT-qPCR (all p-values = 0.05, one-tailed Wilcoxon Rank Sum, n=3) (Figure 2H, J and L).

**Figure 2.**
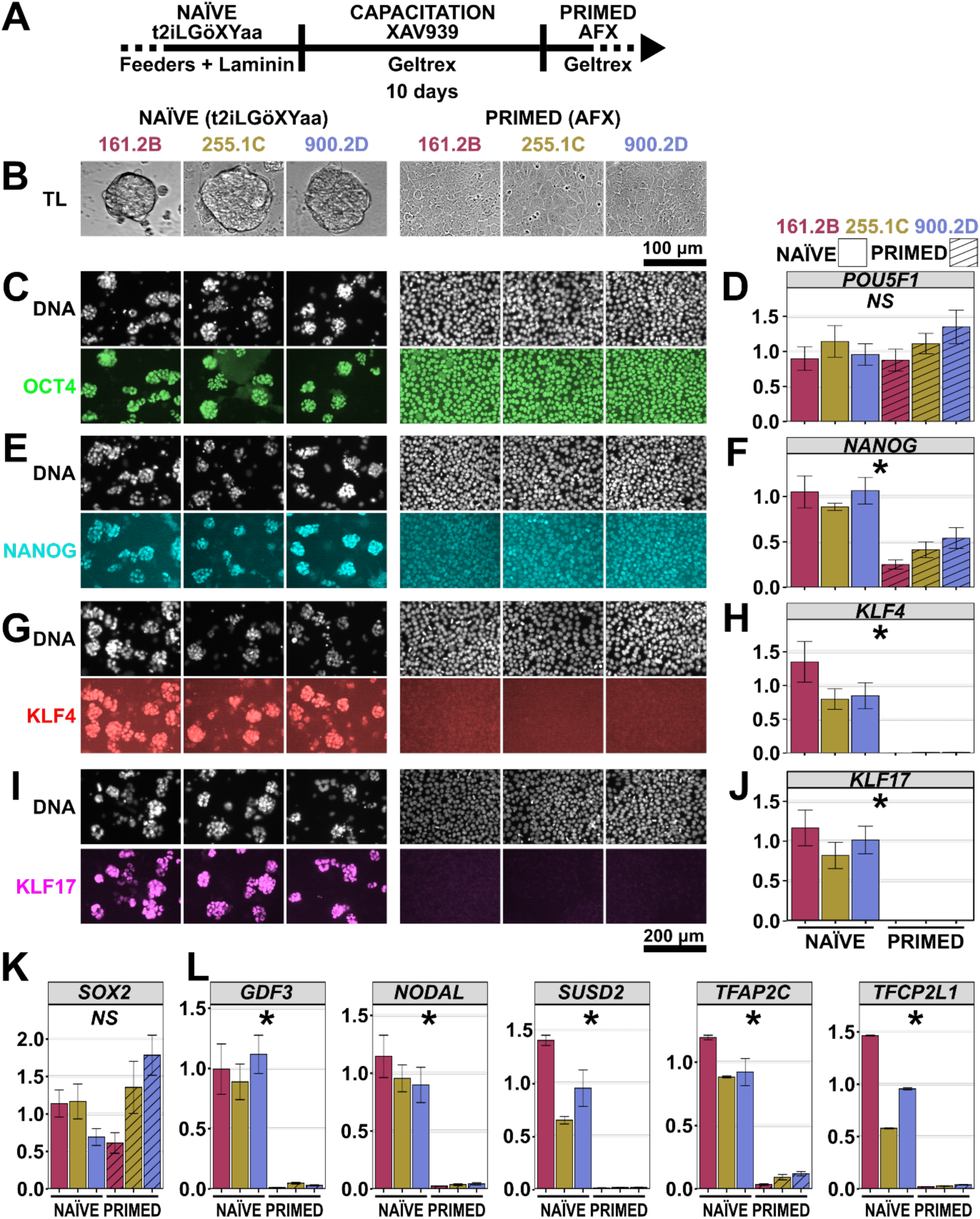
Clonal nPSCs can be capacitated towards the primed pluripotent state. (A) Schematic of nPSC capacitation protocol. (B) Transmitted light images of selected nPSC and corresponding primed PSC lines. (C-J) Immunofluorescence and RT-qPCR of pan-pluripotency markers, OCT4 and NANOG, and naïve pluripotency markers, KLF4 and KLF17 before and after capacitation with XAV939. (K,L) RT-qPCR of pan-pluripotency marker *SOX2*, and naïve markers *GDF3*, *NODAL*, *SUSD2*, *TFAP2C* and *TFCP2L1*. RT-qPCR expression levels are relative to *ACTB*. * denotes a p-value = 0.05 and not significant (NS) denotes p-value > 0.05 for a one-tailed Wilcoxon Rank Sum Test (n = 3).

The naïve signature of clonal nPSCs was investigated through bulk RNA sequencing with inclusion of additional cell lines totalling 13 lines from 5 embryos and 4 donors, as well as published control HNES1, HNES3, and chemically reset (cr)-H9 cell lines (Guo et al., 2016; Thomson et al., 1998). Cells were collected in bulk along with the inactivated murine embryonic fibroblast ‘feeder’ cells (iMEFs), followed by RNA extraction and sequencing, and human reads were separated from mouse reads *in silico* (Figure 3A). In tandem, we sequenced the three capacitated lines from our case study (Figure 2), the control capacitated HNES1 and HNES3 lines, and the embryo derived H9 line. Analysis of these primed lines provided a benchmark against which to evaluate the naïve transcriptome.

**Figure 3.**
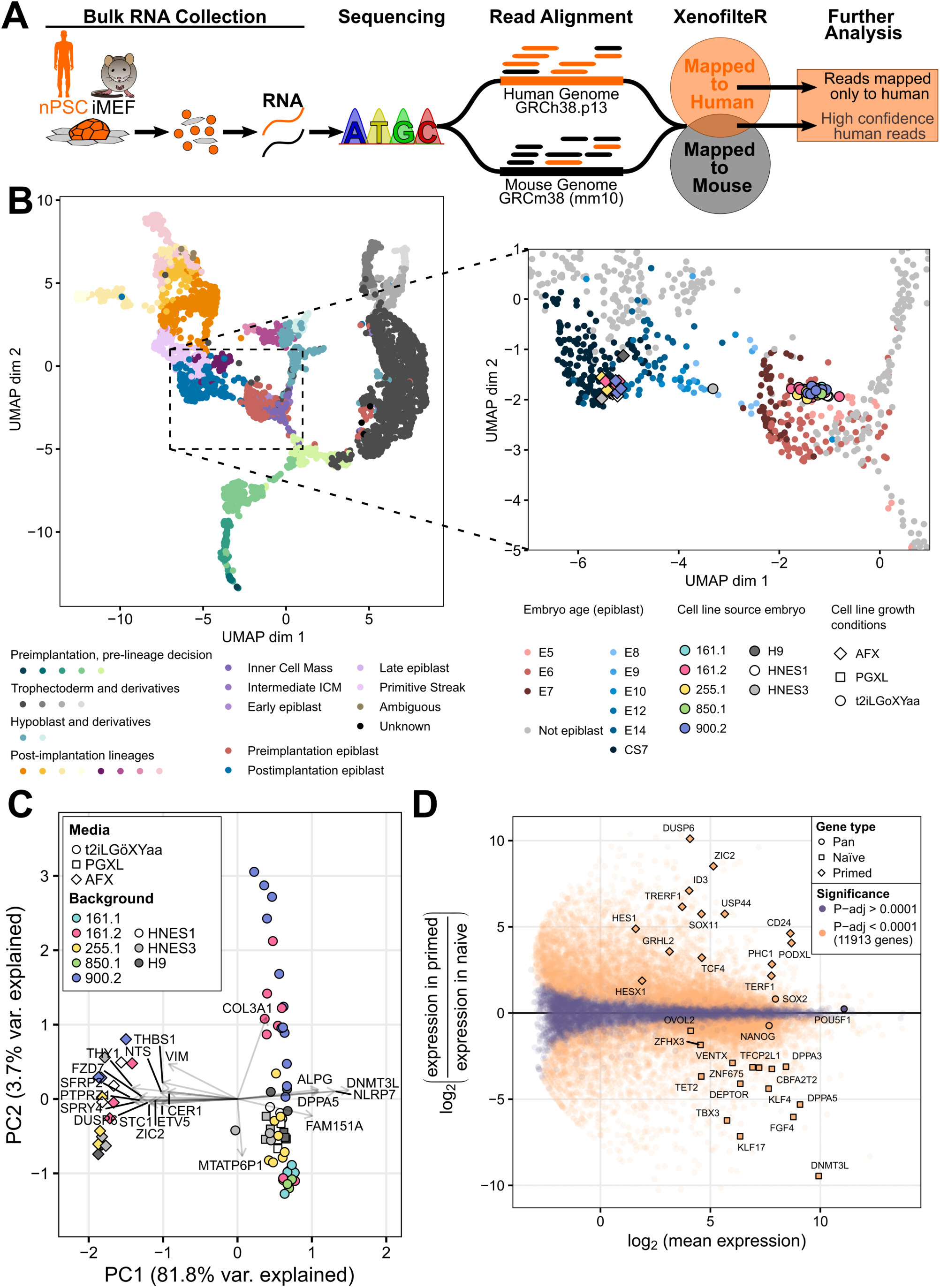
Bulk RNA sequencing reveals high conservation of naïve transcriptomic profile across genetic backgrounds. (A) Schematic of the bulk RNA sequencing pipeline. (B) Left: Single-cell RNA-sequencing reference map of human embryonic populations from (Zhao et al., 2025). Right: Bulk RNA sequencing of our clonal lines and controls mapped onto this reference. Pre- and post-implantation epiblast reference points are color-coded by embryonic time; our samples are color-coded by donor embryo, with growth conditions indicated by shape. Non-peri-implantation reference cells are grey. (C) Principal component analysis (PCA) of autosomal RNA expression in naïve and primed PSCs. Each marker represents an RNA sample from a cell line, with three biological replicates per line. Top 20 genes with the highest coefficient of variation are highlighted. (D) Differential expression analysis between naïve and primed PSCs, with significant (purple) and non-significant (orange) genes. Selected pan-pluripotency, naïve, and primed markers are highlighted.

Using the Early Embryogenesis Prediction Tool (Zhao et al., 2025) we mapped the cell lines to the transcriptional space of the early human embryo, which showed that the naïve clonal and control lines fell within the pre-implantation epiblast cluster near E5/E6 (Carnegie Stage, CS3-2/CS4), and the primed lines within the post-implantation epiblast cluster at E12 (CS7) cells (Figure 3B). Naïve and primed identities were further confirmed by analysing selected gene expression for each of the 13 clonal lines alongside control HNES1, HNES3 and H9 lines (Figure S1A). The chromosomal sex of each line was assessed by expression of the X-linked and Y-linked genes XIST and KDM5D, respectively (Figure S1B) and validated by total X-linked and Y-linked RNA expression (Figure S1C), confirming characterisation of 7 XX lines from 2 genetic backgrounds and 6 XY lines from 3 genetic backgrounds for the clonal lines. X-linked expression in XX lines was less than twice that of XY lines in the naïve state (p-value = 3.5 x 10^-15^, N-way ANOVA) (Figure S1D), which is consistent with X-chromosome dampening described previously (Dror et al., 2024; Petropoulos et al., 2016). Upon capacitation, XX lines displayed lower X-linked expression relative to the naïve state (p-value = 1.4×10^-4^, N-way ANOVA), while XY lines had higher X-linked expression (p-value = 3.9 x 10^-6^, N-way ANOVA), consistent with X-chromosome inactivation in female lines during differentiation. Principal component analysis (PCA) revealed separation by cell state along principal component (PC)1 (76.5% of variance explained) and by sex along PC2 (7.6% of variance explained) (Figure S1E). Sex-based differences could be explained primarily by X- and Y-linked expression; after excluding sex-linked genes, lines did not cluster by sex (Figure S1F) so these genes were excluded from downstream analysis.

PCA of autosomal RNA expression separated the naïve (t2iLGöXYaa and PGXL) and primed (AFX) samples along principal component 1 (PC1) (81.8% of variance explained) (Figure 3C). Loadings displayed expected naïve markers, including *DNMT3L* and *DPPA5*, grouped with the naïve samples, whereas primed samples show grouping with *SPRY4*, *ETV5*, and *DUSP6*, consistent with activity of the FGF/MAPK pathway. In total 11,913 genes were differentially expressed between naïve and primed cells (Figure 3D). Mapping a selected gene panel onto the differential expression plot showed naïve and primed markers sorting to the expected cell type (Adewumi et al., 2007; Blakeley et al., 2015; Guo et al., 2016; Osnato et al., 2021; Stirparo et al., 2018; Xiang et al., 2020). Pan-pluripotency markers *SOX2* and *NANOG* exhibited little difference, with *OCT4* showing no difference between the two cell-states.

We compared control clonal nPSC lines alongside previously established HNES1, HNES3, and cr-H9 in t2iLGöXYaa and a minimal naïve medium, PGXL (Guo et al., 2021) (Figure S2). Similar discrete, domed morphology was observed for colonies in both naïve media (Figure S2A). PCA analysis separated samples by genetic background across PC1 and PC2 (44.8% and 20.2% of variance explained respectively), while PC3 (17.8% of variance explained) primarily separated samples by media condition (Figure S2B). Analysis of contributions of the most variable genes to PC3 failed to show any obvious expression patterns (Table S2). This likely reflects the transcriptomic similarity between cells in the two conditions. Examination of the selected gene panel revealed qualitatively similar expression patterns of pan-, naïve, and primed pluripotency genes (Figure S2C). This is further supported by the differential expression analysis, where only 13 genes were differentially expressed (Figure S2D and Table S3).

### Clonal nPSCs exhibit extra-embryonic lineage potential

Naïve PSCs and epiblast cells isolated from D6 blastocysts have been shown to differentiate into trophoblast (Guo et al., 2021; Io et al., 2021). We therefore challenged our PGXL-adapted case study cell lines (161.2B, 255.1C and 900.2D) to form trophoblast in response to culture in medium supplemented with PD0325901 (PD03; MEK/ERK inhibitor), A83-01 (A83; ACTIVIN/NODAL inhibitor) and Y-27632 (Y; ROCK inhibitor) (PAY). By day 3 all three lines underwent epithelialisation, and by day 7 trophosphere-like cysts emerged from the 161.2B and 255.1C lines (Figure 4A). After 7 days of culture in PAY, all lines produced cells that had down regulated the epiblast marker SOX2 and upregulated the trophoblast marker GATA3 at both protein and RNA levels (Figure 4B and C). Additional RT-qPCR analyses revealed down regulation of pan-pluripotency markers *NANOG* and *OCT4* (*POU5F1*) (Figure 4D), transient *CDX2* expression and upregulation of *GATA2* and *TEAD1*, consistent with progression to trophoblast (Figure 4E).

**Figure 4.**
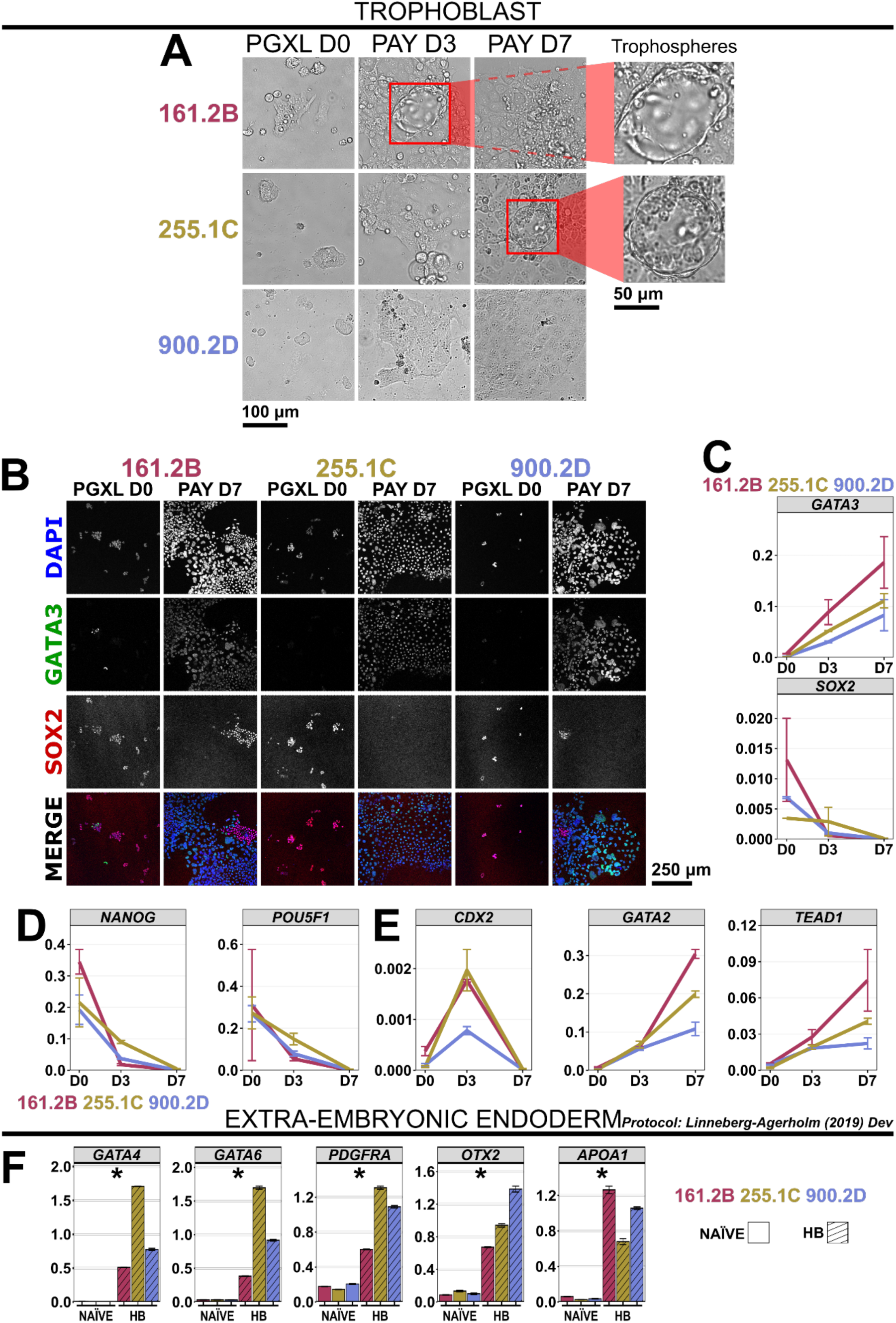
Clonal nPSCs demonstrate extra-embryonic lineage potential. (A) Bright field images of trophoblast differentiation. (B,C) Immunofluorescence images (B) and RT-qPCR (C) of trophoblast marker GATA3 and pan-pluripotency marker SOX2 for trophoblast differentiation of nPSCs. (D) RT-qPCR of pan-pluripotency markers *NANOG* and *POU5F1* (*OCT4*). (E) RT-qPCR of trophoblast markers *CDX2*, *GATA3* and *TEAD1*. (F) RT-qPCR of extraembryonic endoderm markers *GATA4*, *GATA6*, *PDGFRA*, *OTX2* and *APOA1* for hypoblast differentiation of nPSCs, where * denotes a p-value = 0.05 for a one-tailed Wilcoxon Rank Sum Test (n = 3). RT-qPCR levels are relative to *ACTB* for neuroectoderm and paraxial mesoderm and *UBC* for definitive endoderm.

To determine whether our case study lines could also generate extra-embryonic endoderm, the second extra-embryonic tissue produced in the blastocyst, we performed differentiation by means of 7-day induction using LIF, ACTIVIN-A and inhibition of GSK3, via CHIR99021 (Linneberg-Agerholm et al., 2019). RT-qPCR analyses exhibited up-regulation of hypoblast associated markers *GATA4*, *GATA6*, *PDGFRA*, *OTX2*, and *APOA1* in differentiated cells relative to nPSCs (Figure 4F) (all p-values = 0.05, one-tailed Wilcoxon Rank Sum, n=3).

### Capacitated clonal nPSCs exhibit somatic lineage potential

A defining feature of pluripotency is the ability to generate the 3 germ layers: ectoderm, mesoderm, and endoderm. To test this capacity in our case study lines, we performed differentiation to neuroectoderm (NE) (Chambers et al., 2009), paraxial mesoderm (PM) (Loh et al., 2016), and definitive endoderm (DE) (Loh et al., 2014). For all somatic lineage differentiation, clonal nPSCs were capacitated to the primed state to allow efficient differentiation (Rostovskaya et al., 2019). NE differentiation was induced by supplementing N2B27 medium with the ALK3 inhibitor LDN193189 and ALK5 inhibitor A83-01 to supress SMAD signalling (Chambers et al., 2009; Rostovskaya et al., 2019). At the protein level, immunofluorescence revealed that differentiated cells had nuclear expression of NE-associated transcription factors SOX1 and PAX6 (Figure 5A). To verify upregulation of NE markers in differentiated cells relative to the primed PSC starting population, we performed RT-qPCR. As well as testing RNA levels of *SOX1* and *PAX6*, we assessed two additional NE markers, *FOXG1* and *OTX2*. Differentiated cells showed higher RNA levels for all four markers relative to the primed PSCs (all p-values = 0.05, one-tailed Wilcoxon Rank Sum, n=3) (Figure 5B).

**Figure 5.**
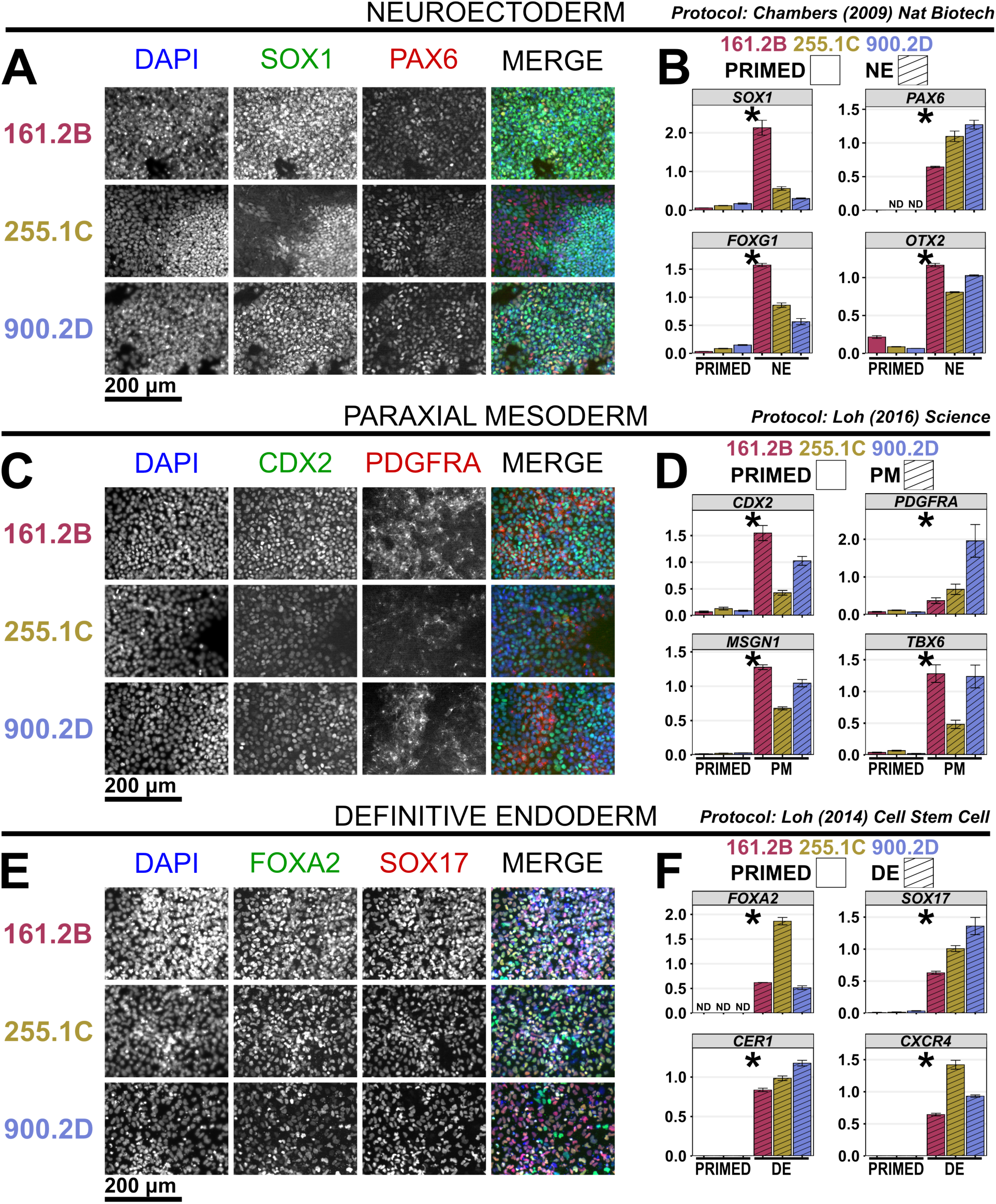
Capacitated clonal nPSCs demonstrate somatic lineage potential. (A) Immunofluorescence images and (B) RT-qPCR of neuroectoderm lineage markers for differentiated nPSCs. (C) Immunofluorescence images and (D) RT-qPCR of paraxial mesoderm lineage markers for differentiated nPSCs. (E) Immunofluorescence images and (F) RT-qPCR of definitive endoderm lineage markers for differentiated nPSCs. RT-qPCR expression levels are relative to *ACTB*. * denotes a p-value = 0.05 for a one-tailed Wilcoxon Rank Sum Test (n = 3).

We next tested the ability of these cell lines to generate PM by initial inhibition of GSK3 and ALK3, followed by FGF induction (Loh et al., 2016). We examined the protein level expression of 2 PM-associated markers: CDX2 and PDGFRA. Differentiated cells showed nuclear expression of transcription factor CDX2 and cell surface expression of receptor PDGFRA (Figure 5C). Acquisition of PM identity was confirmed by RT-qPCR; up-regulation of PM marker RNA was observed for *CDX2*, *PDGFRA*, *MSGN1*, and *TBX6* upon differentiation (all p-values = 0.05, one-tailed Wilcoxon Rank Sum, n=3) (Figure 5D). We challenged our case study cell lines to generate DE as previously published (Loh et al., 2014). Nuclear expression of DE associated transcription factors FOXA2 and SOX17 was observed during DE differentiation (Figure 5E). DE identity was further confirmed by up-regulation in RNA levels of *FOXA2* and *SOX17* and additional DE markers *CER1* and *CXCR4* in differentiated cells (all p-values = 0.05, one-tailed Wilcoxon Rank Sum, n=3) (Figure 5F).

### Analysis of intra-embryo clonal cell lines reveals embryonic mosaicism

To interrogate inter- and intra-embryo RNA expression variation in our 13 selected clonal nPSC lines, PCA was performed using the top 500 variably expressed genes, as determined by coefficient of variation (Figure 6A). Using the first three PCs, individual samples were classified into 5 groups (plus a single outlier which was excluded from further analysis) by unsupervised k-means clustering (Figure 6A, right). Samples from embryo 900.2 were split across groups C4 and C5 (Figure 6A, left). C4 contained samples from 900.2D at passages 15 and 16, while C5 contained the sample at passage 17. Since the later passage sample grouped separately from the earlier ones, the observed expression differences are likely to be a consequence of culture adaptation(s). Differential expression analysis between C4 and C5 resulted in 180 differentially expressed genes that were exclusively up-regulated in C5 (Figure 6B). The directionality of the differentially expressed genes provides further evidence that 900.2A has undergone culture adaptation(s). Differentially expressed genes include the matrix associated components *FN1*, *COL1A1*, *COL1A2*, *KRT24*, and *DCN* and gene ontology pathways analysis of the differentially expressed genes confirmed enrichment of extracellular matrix and cell adhesion proteins, and metalloproteases (Figure 6C). Intra-embryo differences were observed in lines derived from the 161.2 and 255.1 embryos. C1 contained 161.2B while C3 contained 161.2A and 161.2C. Similarly, C1 contained 255.1A while C2 contained 255.1B and 255.1C. In contrast to line 900.2, differential expression analysis gave little insight into the biological relevance of differences between the lines (Figure S3A-C). However, aggregation of the differentially expressed genes between C1 and C2 by chromosome revealed high enrichment on chromosome 5 (Figure S3D), while no such enrichment was observed for differentially expressed genes between C1 and C3 (Figure S3E), suggesting a genetic difference on chromosome 5 between C1 and C2. We therefore explored potential genetic abnormalities within the cell lines.

**Figure 6.**
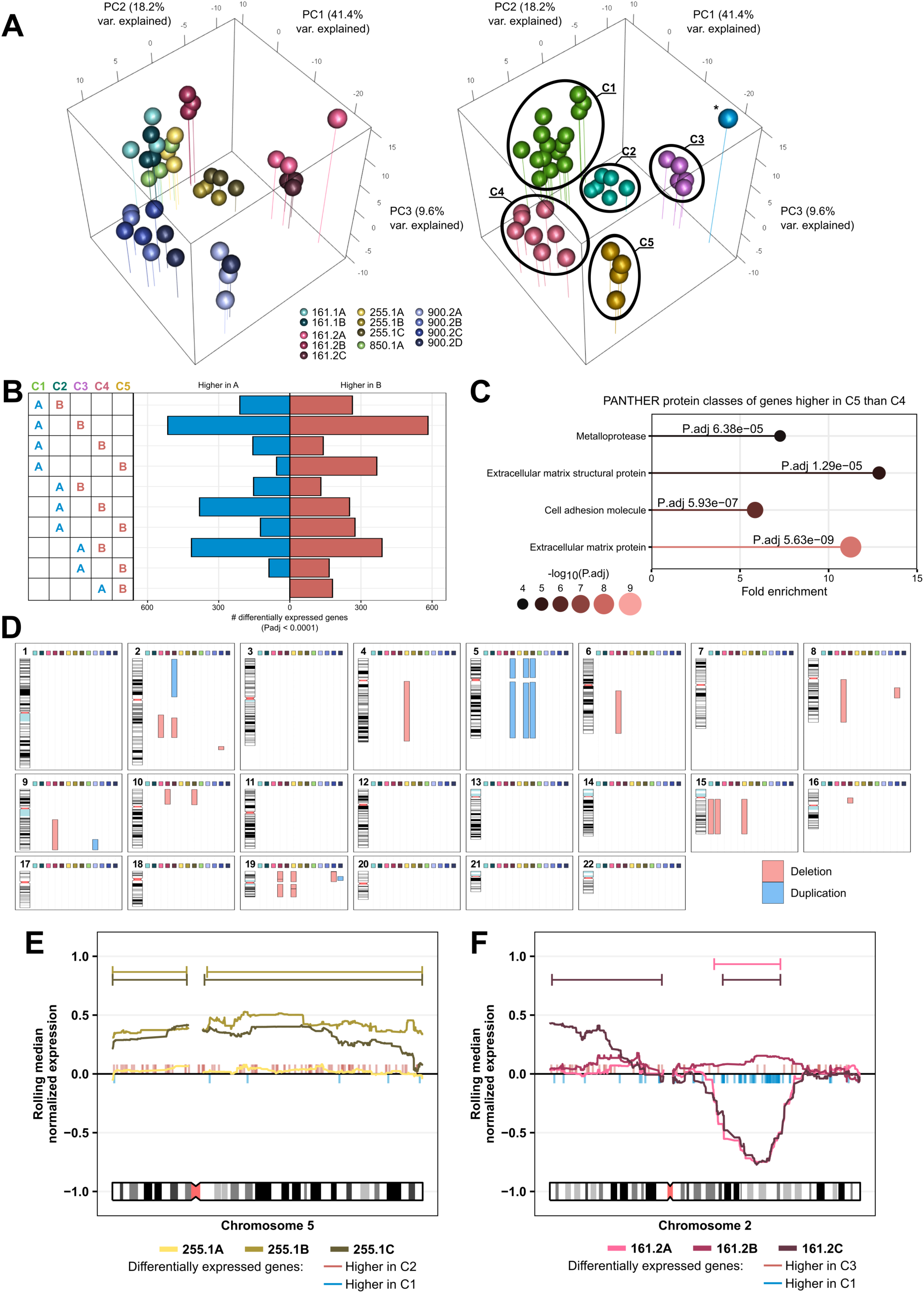
Clonal nPSCs exhibit inter- and intra-embryo variation. (A) Principal component analysis of clonal nPSC lines, with the first three principal components shown. Each marker represents one RNA sample from one cell line and each cell line has three biological replicates from different passages. Left – samples coloured by line. Right – samples coloured by group following unsupervised k-mean clustering. (B) Number of differentially expressed genes (adjusted P value < 0.0001) for different pairwise comparisons of clustered samples. (C) Gene ontology enrichment using PANTHER protein classes for genes that are differentially expressed between clusters 4 and 5. (D) Global mapping of genomic deletions (red) and duplications (blue) across all clonal lines and samples. Chromosomes show banding in black and white, variable regions in blue, and centromeres in red. (E, F) Rolling average RNA expression levels (thick lines) for the indicated chromosomes and cell lines, with differentially expressed genes between relevant clusters indicated with tick-marks and inferred duplication (greater than 0) and deletion (less than 0) events indicated by overhead capped lines.

RNA sequencing read data was used to map segmental duplications and deletions within the genome. This was accomplished by identifying small genetic variants to detect loss of heterozygosity and allelic imbalances, and determining whether these represented gain or loss of genetic material by comparing expression analysis between cell lines (Figure S4). We observed several segmental amplifications and deletions, as well as gain or loss of whole chromosomes and chromosome arms (Figure 6C). In particular, partial and full duplications of Chr5 were seen in multiple lines, consistent with previous reporting in conventional primed PSCs (Halliwell et al., 2021; Ho et al., 2022; Price et al., 2021). We then searched for genetic abnormalities that could explain observed grouping of lines from the 161.2 and 255.1 embryos. For 255.1B and 255.1C lines, duplication-deletion maps (Figure 6D) and rolling average RNA expression maps (Figure 6E) were consistent with a trisomy of Chromosome 5, fitting with the differential expression analysis (Figure S3D); indeed, differentially expressed genes that were higher in 255.1B and C were observed across chromosome 5 (Figure 6E). Interestingly, 161.2A and 161.2C share a segmental deletion in Chr2q that was not observed in 161.2B (Figure 6D, F). Importantly, because this deletion appears in two lines out of three, we do not suspect this to be a culture acquired mutation. Rather, it most likely indicates *in vitro* capture of embryonic mosaicism. We therefore attribute the segregation of clusters 1-3 in transcriptional space to these genetic differences. Unlike the Chr2q deletion in the 161.2 lines, Chr5 trisomy is a common culture acquired mutation; therefore, it is unclear if the genetic difference between the 255.1 lines represents a *bona fide* capture of embryonic mosaicism or independent acquisition of an additional copy of Chr5 during culture.

## DISCUSSION

Derivation of multiple pluripotent stem cell lines per human blastocyst provides a valuable resource for modelling mosaicism and investigating the effects of genetic background and chromosomal anomalies on differentiation potential and lineage regulation during development. We adapted our previous strategy for capturing naïve pluripotent stem cells following immunosurgery and ICM disaggregation from human blastocysts (Guo et al., 2016; Strawbridge et al., 2022) to generate individual pluripotent stem cell lines as a resource (Figure 1), serving as ‘proof-of-principle’ methodology for future studies into intra- and inter-embryo karyotypic variation. Preliminary analysis of the new lines reveals several cases of chromosomal divergence within an embryo. It is hoped that other researchers will be inspired to supplement the stem cell resource by deriving additional clonal lines of assorted genetic backgrounds that will enrich the diversity of the available material and exhibit supplementary mutations to promote further understanding of developmental and post-natal human disorders.

A high burden of chromosomal defects in human embryos generated by *in vitro* fertilisation is implicated from trophectoderm biopsies taken prior to transfer (Yang et al., 2024); around 30% of morphologically normal embryos generated through in vitro fertilization (IVF) have been reported to be mosaic (Buldo-Licciardi et al., 2023). However, more embryos appear competent to develop to term than may be predicted from the trophectoderm analysis (Capalbo et al., 2021). It has been hypothesised that defective cells may either be extruded into the trophectoderm or more frequently arise *de novo* in that tissue (Buldo-Licciardi et al., 2023; Capalbo et al., 2021). Experimental induction of aneuploidies in developing mouse embryos revealed that defective cells within the ICM tend to be eliminated by apoptosis, whereas those within the trophectoderm are less urgently excluded (Bolton et al., 2016). Similar experiments have not been performed using human embryos. Given the increased flexibility of the human epiblast to differentiate into trophectoderm compared with mouse (Guo et al., 2021), it may be considered more likely that defective cells within the human blastocyst are more readily extruded into the trophectoderm lineage. However, current evidence does not support incorporation of ICM cells into the overlying trophectoderm in the intact blastocyst (Corujo-Simon et al., 2024). In short, some uncertainty persists regarding the effective use of trophectoderm or polar body biopsy as an accurate means to identify viable embryos (Dahdouh, 2021; Dahdouh et al., 2023) as the mechanisms associated with persistence or elimination of aberrant cells remain largely unknown. Our clonal nPSCs will present an accessible resource with which to generate blastoids to model human preimplantation development and investigate the effect of genetic aberrations. As well as comparing potency of each line to form the three founder lineages in 2D differentiation studies, chimeric blastoids could be assembled from two karyotypically diverse, differentially labelled lines from the same embryo to monitor the fate of cells bearing known chromosomal aberrations. An advantage of using cell lines from the same embryo is minimisation of elimination of ‘loser’ cells by apoptosis instructed by the genetically distinct ‘winner’ cells via cell competition (Price et al., 2021).

Like the control HNES1, HNES3 and cr-H9 cell lines, our randomly selected case study clonal examples could progress towards the primed state of pluripotency (Figure 2). X and Y-linked gene expression profiles were faithful to the chromosomal sex of each cell line; female lines exhibited levels of X chromosome expression consistent with previously reported chromosome dampening behaviour (Figure S1) (Adewumi et al., 2007; Dror et al., 2024). During capacitation and priming they also appeared to undergo inactivation of one X chromosome, as expected. The clonal lines exhibited some variability in expression levels, as anticipated from their genetic backgrounds, but acquisition of epithelial morphology and basic transcriptional profile characteristic of priming was observed in all lines. Both the naïve and primed states of the new cell lines matched the equivalent expression profiles of pluripotent cells within the embryo (Zhao et al., 2025) (Figure 3). The naïve lines tested were all competent for extra-embryonic differentiation, but variability was observed between the samples, both phenotypically and by RT-PCR (Figure 4). Differentiation from the primed state to each of the three germ layers was achieved (Figure 5). In future, it will be interesting to examine effects of chromosomal anomalies in specific tissues via co-culture of different euploid and genetically aberrant lines via directed differentiation or in chimeric gastruloids.

In-depth profiling of the clonal lines by RNA-sequencing revealed transcriptional and genetic variation between clones of different genetic backgrounds and those derived from single embryos (Figure 6). We speculate that some of these differences are captured directly from embryos as a result of the normal genetic variation between individuals as well as embryonic mosaicism from *de novo* mutations, while other differences may be the result of culture adaptation. Therefore, we anticipate that these lines, as well as future lines derived using these methods, will be a valuable resource for disentangling the relative impacts of genetic variation and non-genetic culture adaptation on the variable behaviour, such as different efficiency in directed differentiation protocols, typically observed with human pluripotent cell lines.

In contrast to the naïve pluripotent stem cell lines derived from mouse embryos, human naïve pluripotent stem cells possess the capacity to give rise to extraembryonic lineages (Guo et al., 2021; Io et al., 2021; Linneberg-Agerholm et al., 2019). This property has been particularly advantageous for the development of 3D organoids resembling human blastocysts, known as ‘blastoids’ (Kagawa et al., 2022; Liu et al., 2021; Yanagida et al., 2021). It is possible that aneuploid cells are eliminated from the epiblast through segregation into the trophectoderm, thus protecting the prospective embryo. Availability of genetically normal and diverse clonal nPSCs could ultimately enable this hypothesis to be tested using blastoids, thereby assisting future interpretation of persistence versus elimination of cells exhibiting adverse properties.

## LIMITATIONS OF THE STUDY

1. The proportion of embryos producing cell lines in this study is quite small; only 10 of 84 morphologically normal embryos resulted in one or more stable cell lines. As expected, a proportion of the plated ICM cells gave rise to flat colonies with morphology consistent with extra-embryonic endoderm or trophoblast identity. Such cells would be expected to have segregated from the epiblast by the time the ICM was disaggregated. Some of the dome-shaped colonies emerging from the initial plating could be passaged a few times, but subsequently failed to thrive. The reason for this could not be decisively determined, but such colonies may have carried or acquired mutations incompatible with self-renewal. Alternatively, it is possible that they may have been casualties of sub-optimal culture conditions.
2. All our cell lines were derived using t2iLGöYaa with Geltrex added at the time of plating. We did not attempt derivation using alternative published culture regimes established elsewhere for expansion of nPSCs, since we had previously successfully derived robust cell lines using our original system.
3. Although transcriptomic profiles were consistent with naïve and primed pluripotent states across the lines, we did not systematically assess differentiation capacity across all clonal lines at this stage, nor did we evaluate multiple clones from the same embryo in parallel.
4. Although we observe highly similar chromosomal abnormalities in two of three clonal lines from the same embryo, we cannot conclusively determine whether these represent bona fide capture of embryonic mosaicism. Our inference relies on the low probability of independent acquisition of the same aneuploidy during culture, but direct evidence from the embryo of origin is lacking.
5. The embryos used for this work were donated with informed consent from couples who had completed their fertility treatment, and were therefore unlikely to be of the highest quality.

## MATERIALS AND METHODS

### Embryo Culture

Supernumerary frozen human embryos were anonymously donated by couples undergoing *in vitro* fertility treatment with informed consent and access to counselling. Use of human embryos in this research is approved by the Multi-Centre Research Ethics Committee, approval O4/MRE03/44, Integrated Research Application System (IRAS) (21/ PR/1231), and licensed by the Human Embryology and Fertilization Authority of the United Kingdom, research license R0178.

Frozen blastocysts were thawed using commercial kits specified by the clinic from which they were donated and cultured in N2B27 medium under mineral oil in a humidified incubator at 37°C, 7% CO2 and 5% O2 for 24 hours before derivation.

Derivations were performed on irradiated mouse embryonic fibroblast (iMEFs) feeder cells in N2B27 medium supplemented with inhibitors of GSK3 (1 μM CHIR99021), MEK/ERK (1 μM PD0325901), aPKC (2.5 μM Gö6976), ROCK (10 μM Y-27632) and the addition of LIF (10 ng/mL) and ascorbic acid (250 μM) (t2iLGöYaa), with a laminin spike-in following seeding (5 μL of 1 mg/mL). From passage 4 onward, the medium was modified by reducing CHIR99021 to 0.5 μM, reducing ascorbic acid to 125 μM, and adding Tankyrase inhibitor (2 μM XAV939), resulting in the final composition referred to as t2iLGöYXaa. After lines were sufficiently expanded, the laminin spike-in was replaced with dilute Geltrex (20 μL of 1:4 in DMEM/F12). Please refer to (Strawbridge et al., 2022) for full protocol.

### Pluripotent stem cell (PSC) maintenance

Cells were cultured throughout in a humidified incubator with 5% O2 and 7% CO2 at 37°C. Established nPSCs were maintained on iMEFs in t2iLGöYXaa or PGXL (1 µM PD0325901, 2 µM Gö69832, 2 µM XAV939 10 ng/ml human LIF in N2B27). Inhibition of ROCK (10 μM Y-27632) was performed during plating of cells in PGXL. Both t2iLGöYXaa and PGXL conditions are accompanied by a Geltrex spike-in. Cells were passaged using Accutase. Conventional primed PSCs were cultured in N2B27 supplemented with ACTIVIN-A (20 ng/mL), FGF2 (12.5 ng/mL), and XAV939 (2 µM) (AFX) on Geltrex pre-coated plates (1:100 in DMEM/F12, 1 h at 37°C) and passaged using 0.5 mM EDTA in PBS.

### PSC differentiation

PSCs were capacitated from the naïve to the primed state following the protocol previously described (Rostovskaya et al., 2019). Briefly, naïve ES cells were plate for 48 hours on geltrex-coated plates in PGXL+Y. The next day, cells were washed once with PBS and the medium was switched to N2B27+ 2 μM XAV939 for up to 10 days. For trophoblast differentiation, naïve (n)PSCs cultured in PGXL on mitotically inactivated mouse embryonic fibroblasts (MEFs) were passaged onto Geltrex-coated plates in PGXL supplemented with 10 µM Y-27632 to remove feeder cells. When confluent, cultures were replated once more onto Geltrex in PGXL+Y. The following day, cells were washed and transferred to PAY medium, consisting of 1.5 µM PD0325901, 1 µM A83-01, and 10 µM Y-27632 in N2B27, where they were maintained for five days to promote differentiation. Hypoblast differentiation was carried out according to the method previously established (Linneberg-Agerholm et al., 2019). To induce differentiation into somatic lineages, established protocols were followed for each germ layer. Neuroectodermal differentiation was performed using the approach outlined previously (Chambers et al., 2009), while paraxial mesoderm differentiation was conducted as previously described (Loh et al., 2016). For definitive endoderm differentiation, cells were treated following according to the published protocol (Loh et al., 2014).

### Immunocytochemistry

Whole embryos were fixed in 4% paraformaldehyde (PFA) for 15 minutes at room temperature (RT), followed by a PBS wash containing 3 mg/ml polyvinylpyrrolidone (PBS/PVP). Samples were permeabilized in 0.25% Triton X-100 in PBS/PVP for 30 minutes and then blocked in PBS containing 2% donkey serum, 0.1% bovine serum albumin (BSA), and 0.01% Tween 20 for 1 to 2 hours at RT. Primary antibodies were diluted 1:200 in blocking buffer and incubated overnight at 4°C. The next day, samples were incubated with secondary antibodies (diluted 1:500 in blocking buffer) for 1 to 2 hours at RT in the dark. Both antibody incubations were followed by three 15-minute washes in blocking buffer.

Cells were fixed in 4% PFA for 20 minutes at RT and then washed 3 times with PBS. Samples were and permeabilized in PBS containing 0.5% Triton X-100 for 30 minutes and blocked in blocking buffer (5% BSA, 5% donkey serum, 0.25% Triton-X in PBS) for 30 minutes. Primary antibody incubation was performed overnight at 4°C in blocking buffer (see Table S4 for list of primary antibodies), followed by three 15-minute washes in 0.02% Triton X-100. Samples were incubated with secondary antibodies in blocking buffer for 1 hour at RT in the dark then washed three times in PBS.

### RNA purification

To prepare RNA for sequencing, cells were dissociated using accutase, diluted in DMEM/F12 and centrifuged to pellet the cells. Supernatant was aspirated and the resulting pellet was flash frozen on dry ice and stored at −70°C until all samples were collected. RNA was collection using TRIzol (ThermoFisher 15596026) followed by column purification with the PureLink RNA mini kit (ThermoFisher 12183020) In brief, pellets 1mL Trizol added and pipetted up and down and transferred to a 1.5ml Eppendorf tube. Chloroform (20μl) was added and the tubes incubated at RT for 2-3min. The samples were centrifuged for 15min at 12000g at 4°C. We kept both the upper aqueous phase containing RNA, and the lower phenol phase, containing the protein. To the aqueous phase, equal volumes of 70% ethanol and mixed by vortexing. This was then purified using the PureLink RNA mini kit, following the kit instructions. The sample was bound to the column, and oncolumn DNase treatment was carried out PureLink DNase set (12185010) to remove any contaminating genomic DNA. After washing and drying the column, the RNA was eluted with 2 x 50μl RNase-free water, and the samples were stored at −70°C.

For RT-qPCR, RNA was extracted using a Monarch Total RNA Miniprep kit according to the manufacturer’s instructions with on-column DNase treatment.

### Real time quantitative-PCR

cDNA was prepared from total RNA using RT-qPCR was performed using SuperScript IV First-Strand Synthesis System with oligo d(T)20 primers according to the manufacturer’s instructions. RT-qPCR was performed PowerUp SYBR Green Master Mix for qPCR according to the manufacturers’ instructions on a QuantStudio 12K Flex using target-specific primers (see Table S5).

### Bulk-RNA sequencing and read alignment

Initial quality control of the RNA was performed with Agilent RNA tapestation reagents (5067- 5576; 5067-5577; 5067-5578) and the Qubit RNA HS Assay Kit (Q32855). Sample library production was performed using 500ng of total RNA with the NEBNext Ultra II Directional RNA Library Prep Kit for Illumina (E7760) and the NEBNext Multiplex Oligos for Illumina (E6440), with 96 unique dual index primer pairs, in conjunction with NEBNext Poly(A) mRNA 15 Magnetic Isolation Module (E7490). This was performed according to manufacturer’s instructions. Quality control of the resulting libraries was performed with Qubit dsDNA HS Assay Kit (Q32854) and Agilent DNA 5000 tapestation reagents (5067-5588; 5067-5589). Samples were then pooled in equimolar quantities and sequenced on a Novaseq 6000 on two lanes of a 1 flowcell as paired end 150bp (PE150) reads.

Barcodes and low-quality bases were trimmed from reads using TrimGalore V0.4.1 as a wrapper for Cutadapt V1.8.1. Trimmed reads were independently aligned to the GRCh38.p13 human genome and annotation, and the GRCm38.p6 (mm10) mouse genome and annotation using STAR V2.7.6a with per-sample 2-pass mapping. The flag “-outSAMattributes NM” was used for compatibility with XenofilteR. Mouse reads were filtered out from the human alignment by passing both sets of aligned reads to XenofilteR V1.6, with MM-threshold set to 8. Raw counts were generated using the featureCounts function of subread V1.5.1.

### Mapping to human embryonic reference

Sequencing data were mapped to the human embryonic reference using the Early Embryogenesis Prediction tool V2.1.1 (Zhao et al., 2025) and visualized using R V4.1.1.

### Gene expression analysis

Gene expression analysis was performed using R V4.1.1. Counts per million were generated using normalized library sizes with the cpm function from edgeR V3.36.0. Unless otherwise specified, genes on the X and Y chromosomes were excluded from analysis to reduce sex-dependent differences between samples. PCA analysis was performed using the 500 genes with the greatest coefficient of variance across relevant samples. Differential gene expression analysis was performed using edgeR.

### Gene ontology enrichment analysis

Gene ontology enrichment analysis was performed using the PANTHER overrepresentation test using PANTHER version 19 and the PANTHER protein class annotation set. A Fisher’s Exact test was used with False Discovery Rate correction for multiple testing within a single comparison. No further correction was applied for multiple comparisons.

### Chromosomal duplication and deletion analyses

Bulk RNA-sequencing data was used to map putative deletions and duplications in lines. Reads were realigned to mouse genome GRCm39 for improved accuracy, using the same methods described above. After filtering for human reads with XenofilteR, a database of all short variants from samples grown in t2iLGoXYAA were generated using GATK V4.0.10.1 tools. Briefly, read groups were added using AddOrReplaceReadGroups, reads were split with SplitNCigarReads. Quality scores were recalibrated using BaseRecalibrator with known sites of variants taken from the dbsnp138 reference downloaded from the GATK public resources database, and recalibrations were applied using ApplyBQSR. Variants were called with HaplotypeCaller, outputting GVCF files. These were aggregated into a database with GenomicsDBImport, and genotypes were called with GenotypeGVCFs. Variants were exported using VariantsToTable. Subsequent analysis was performed in R V4.1.1. Analysis of genetic anomalies was split into two phases; detection of loss of heterozygosity to identify deletions with full penetrance in clonal lines, and analysis of biased allelic expression followed by differential expression analysis to detect duplications and subclonal deletions. For loss of heterozygosity analysis, filtered variants were called as heterozygous or homozygous. Each chromosome arm was segmented into regions with consistent het/hom variant ratios using circular binary segmentation (via the CNA function from DNAcopy V1.68.0) on the binarized data to detect change points. All segments within a sample were then tested for a decrease in homozygosity with a one-tailed binomial test, p-values were corrected with Holm multiple testing correction, and segments with an adjusted p-value < 1×10^9 and a lower 95% confidence interval of 90% homozygous were considered genuine regions of loss of heterozygosity. Next, contiguous regions of chromosome arms that did not display loss of heterozygosity were segmented into regions with consistent allelic expression ratios. For each heterozygous variant, the major allelic ratio (higher expressed allele divided by lower expressed allele) was calculated, and circular binary segmentation (via the CNA function from DNAcopy) was performed on this ratio to detect change points (Olshen et al. 2004; Venkatraman and Olshen 2007). All segments within a sample were then tested for an increase in the major allelic ratio (indicative of an allelic imbalance) using a non-parametric one-tailed Brunner-Munzel test (using the brunner.munzel.test function of lawstat V3.6). P-values were corrected with Holm multiple testing correction, and segments with an adjusted p-value less than 0.01 were considered genuine regions with allelic imbalance. Finally, expression analysis was used to define segments with allelic imbalance as duplications, deletions, or undetermined. Genes were catagorised as expressed or non-expressed by taking the median expression (in counts per million, generated with normalized library sizes using the cpm function of edgeR) across all samples and performing Gaussian mixture modelling with two components. RPKM values were generated for expressed genes by dividing counts per million by gene length in kilobases. These values were log transformed and centred across all samples. Segments were considered to have a significant change in expression if the centred, log transformed expression values for genes contained within them significantly differed from 0 (two-sided t-test, P-value < 0.01); segments with significantly reduced expression were called as deletions while segments with significantly increased expression were called as amplifications.

## ACKNOWLEDGEMENTS

We are indebted to the clinical embryologists, particularly Karen Thompson and Catherine Pretty, the nurses and patients of the assisted conception clinics for provision of human embryos left over from fertility treatment. We thank Austin Smith and his team for willingness to bank and distribute the cell lines.

## AUTHOR CONTRIBUTIONS

Original conceptualization and funding acquisition: J.N.; patient recruitment and provision of human embryos: K.T. and C.P.; lab-based optimisation of methodology and instigation: J.N. and G.G.; expansion and banking of cell lines S.E.S and K.A.J.; investigation, S.E.S., L.E.B., C.R., K.A.J., T.A., T.L., M.P., V.M., A.L.C. and J.N.; analytical methodology, S.E.S., L.E.B., M.R. and J.C.; resources, K.T., C.P. and J.N.; data curation, S.E.S. and L.E.B.; software, L.E.B.; supervision, J.N., S.E.S., L.E.B.; writing, reviewing and editing: all authors.

## DATA AVAILABILITY

Code used for RNA-sequencing data analysis can be found at https://github.com/L-E-Bates/clonal_hnPSC_derivation RNA sequencing data have been deposited in the BioStudies database (https://) under accession number E-MTAB-15169.

---

**Figure S1.**
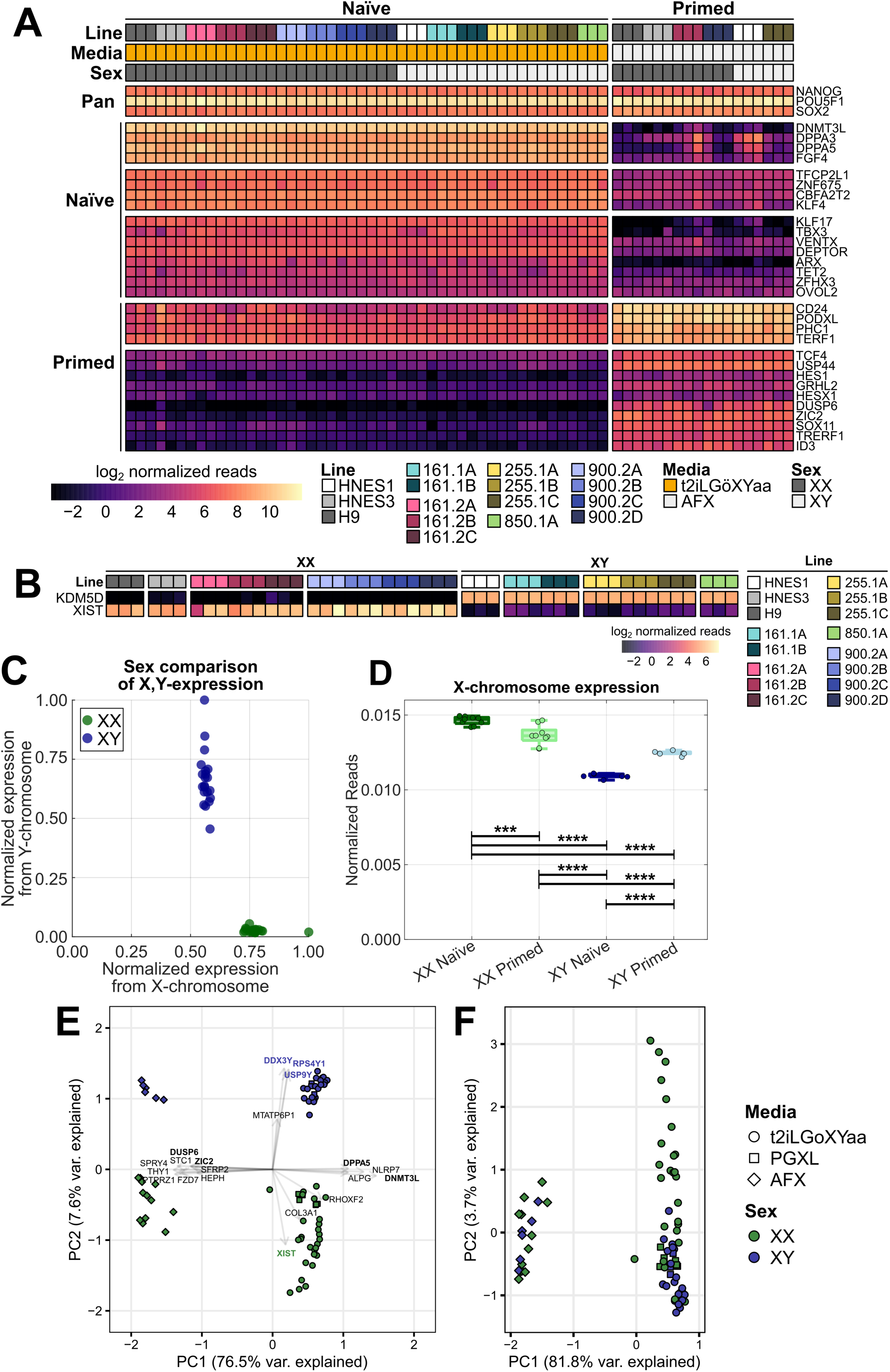
Analysis of bulk-RNA sequencing and sex chromosome regulation. (A) Heatmap of the selected pan, naïve and primed pluripotency associated genes in replicates of clonal and control lines. (B) Heatmap of Y-linked gene KDM5D and X-linked gene XIST. (C) Total normalized X- and Y-linked expression for all clonal lines, coloured by sex. (D) Boxplot summarizing total normalized X-linked reads for XX and XY samples in naïve or primed culture conditions (*** P<0.001, **** P<0.0001, N-way ANOVA with Tukey-Kramer post-hoc correction). (E) Principal component analysis of the 500 genes with the greatest coefficient of variation from clonal and control lines in naïve and primed culture conditions. (F) Principal component analysis of the 500 genes with greatest coefficient of variation after excluding X- and Y-linked genes as in Figure 3C.

**Figure S2.**
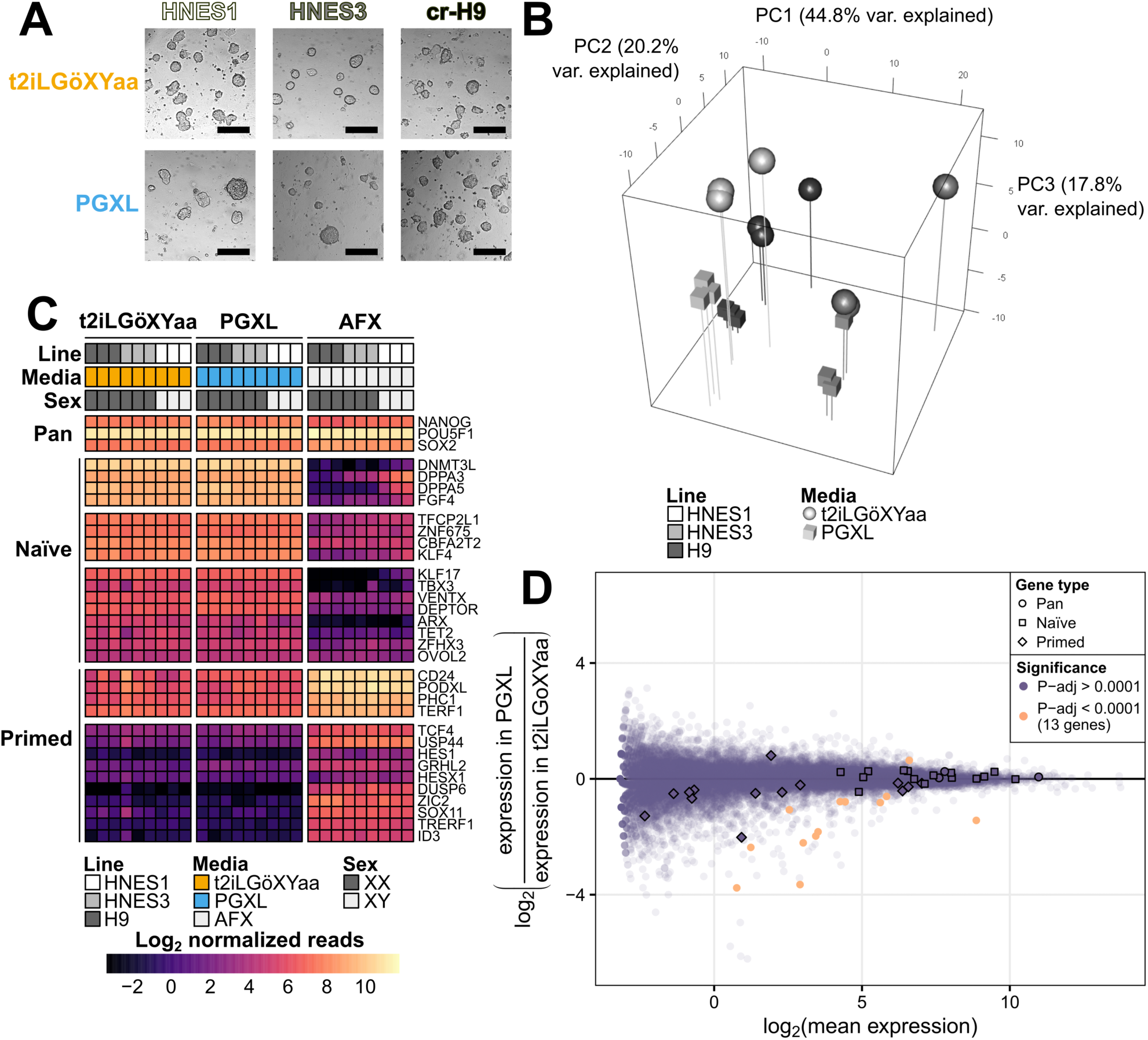
Minor transcriptomic variation detected between nPSC culture conditions. (A) Bright field images of established nPSC lines in derivation medium (t2iLGöXYaa) and a minimal medium (PGXL). (B) Principal component analysis of autosomal RNA expression for different nPSCs culture conditions. (C) Heatmap of selected pan-, naïve, and primed pluripotent genes. Expression values are log2(CPM). (D) Differential expression analysis between nPSC culture conditions.

**Figure S3.**
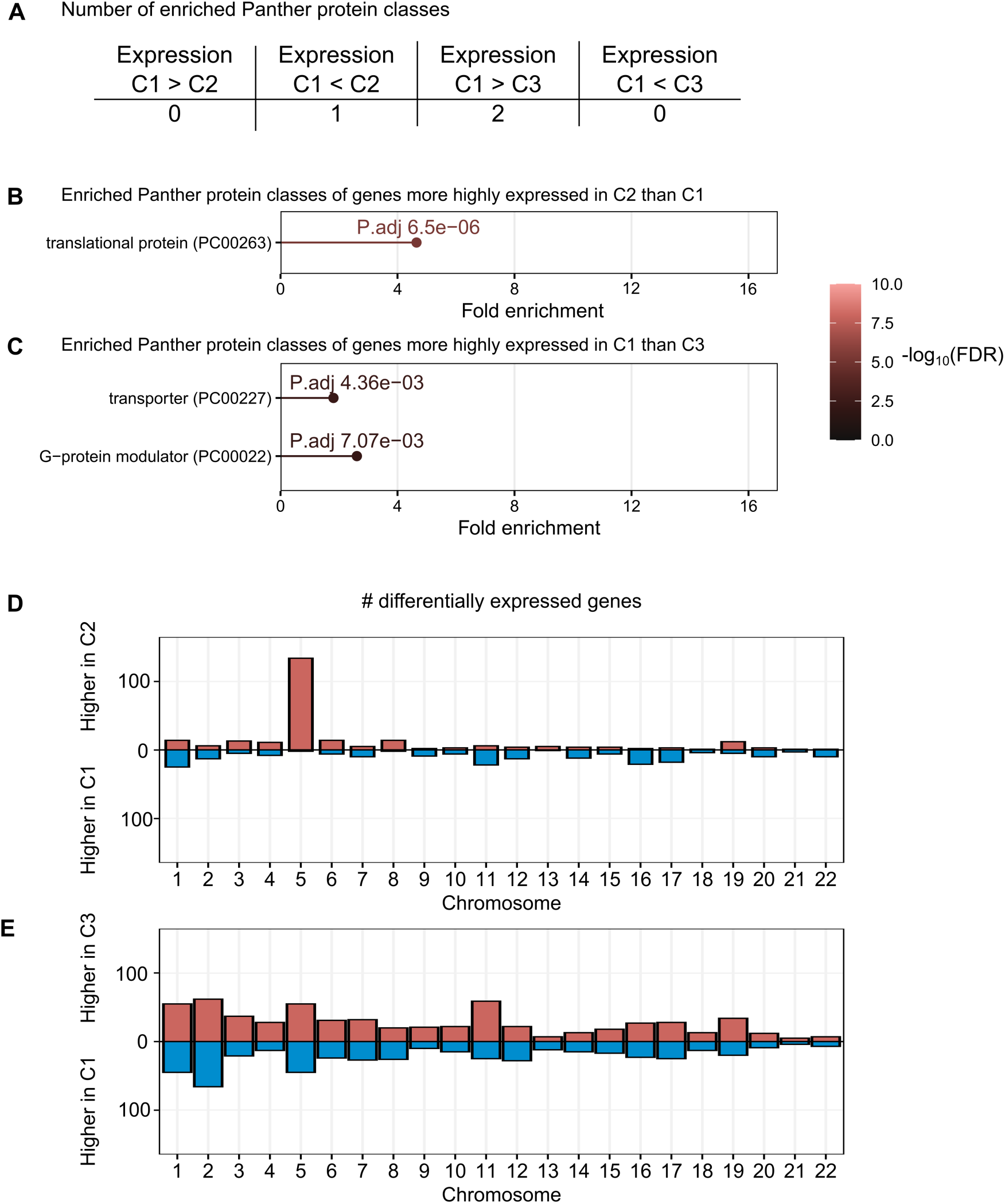
Differential gene expression analysis of cluster 1 versus clusters 2 and 3. (A) Summary of the number of enriched protein classes among differentially expressed genes between cluster 1 and clusters 2 and 3. (B,C) Enriched protein classes among differentially expressed genes. (D, E) Differentially expressed genes between cluster 1 and cluster 2 (D) and cluster 3 (E) aggregated by chromosome.

**Figure S4.**
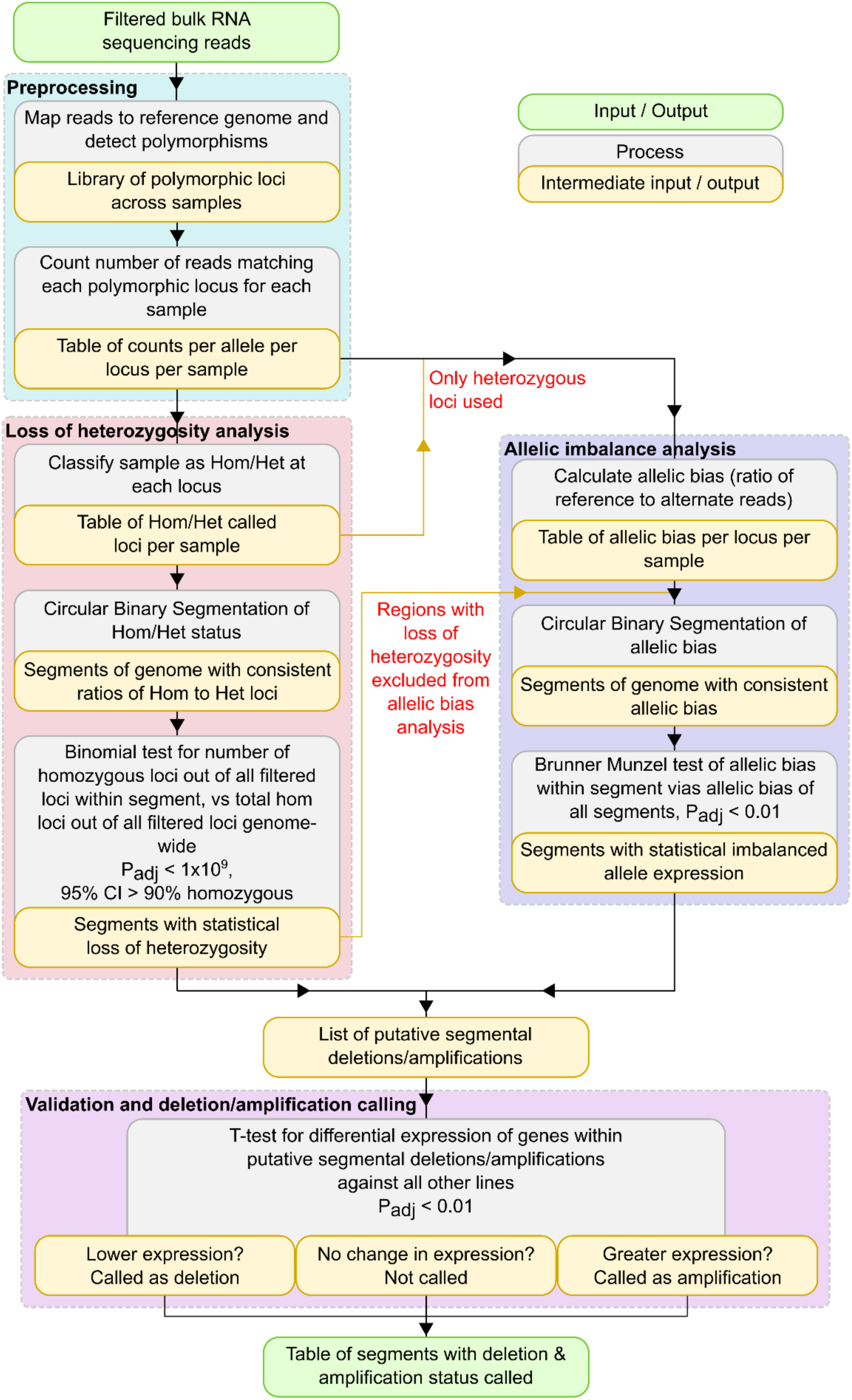
Duplication-deletion analysis pipeline. Flowchart of the pipeline used to infer deletions and amplifications from bulk RNA sequencing data. The analysis uses polymorphisms to detect for each sample regions with loss of heterozygosity (such as regions in which only one allele is present due to a segmental deletion) and regions with allelic imbalance (such as regions with a segmental duplication increasing the copy number of one allele relative to the other, or regions carrying deletions in only a fraction of the population). Once these regions have been identified, they are validated and called as deletions versus amplifications by identifying differences in gene expression across the region relative to all of the other samples.

**Table S1.**
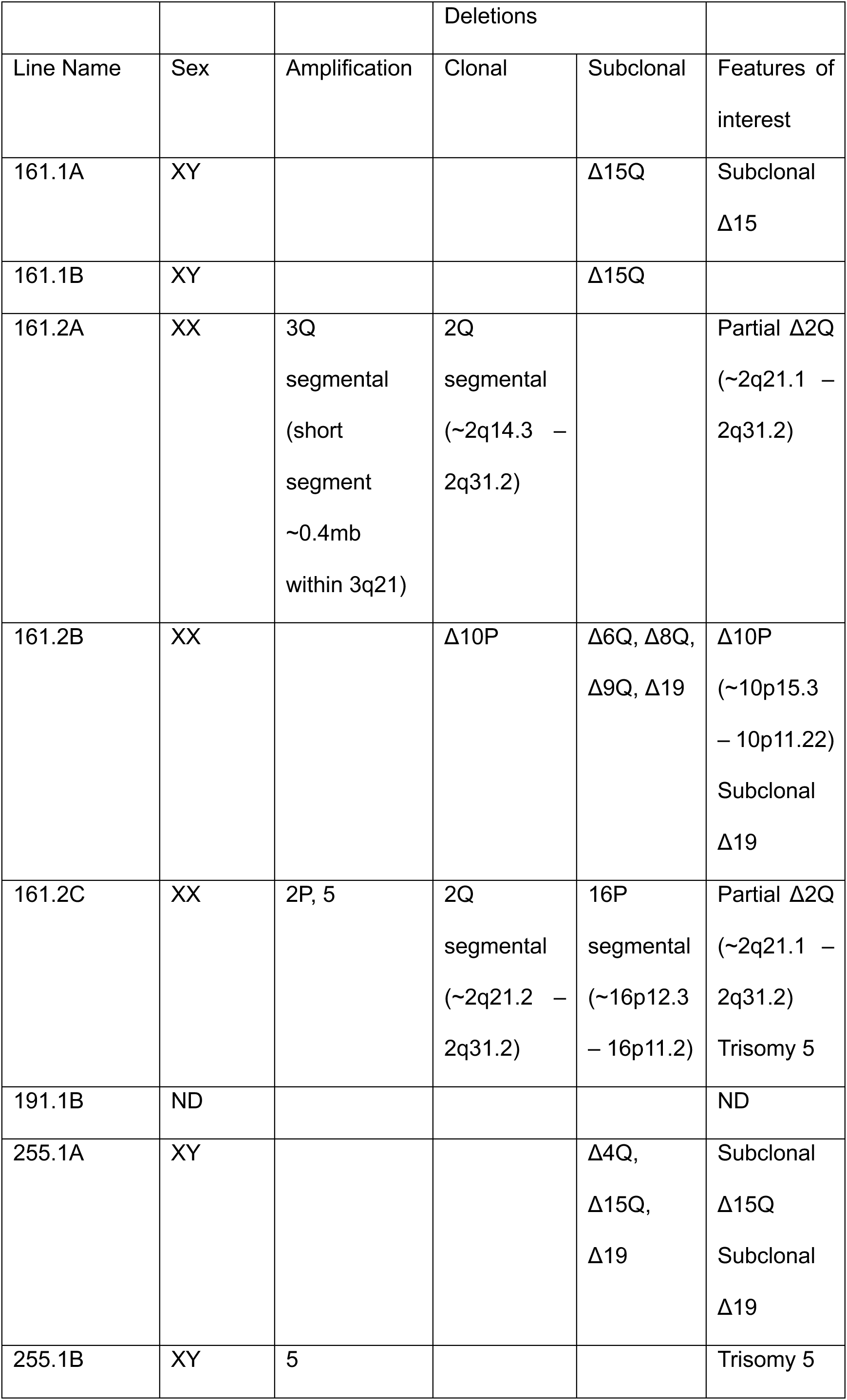
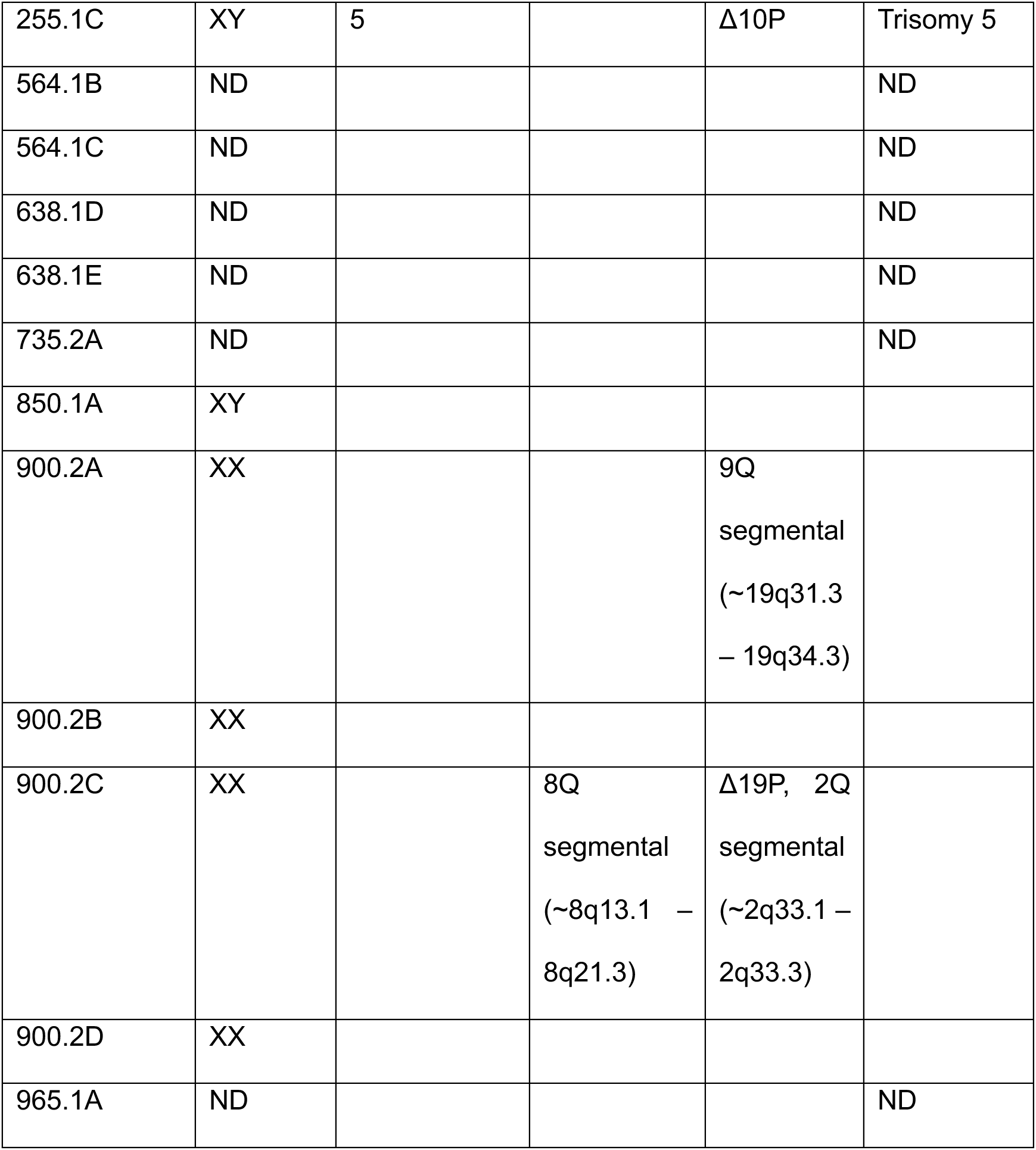
Summary of clonal naïve pluripotent cell lines derived Chromosomal sex and summary of genetic anomalies detected in the clonal cell lines derived in this work. ND = not determined.

**Table S2.**
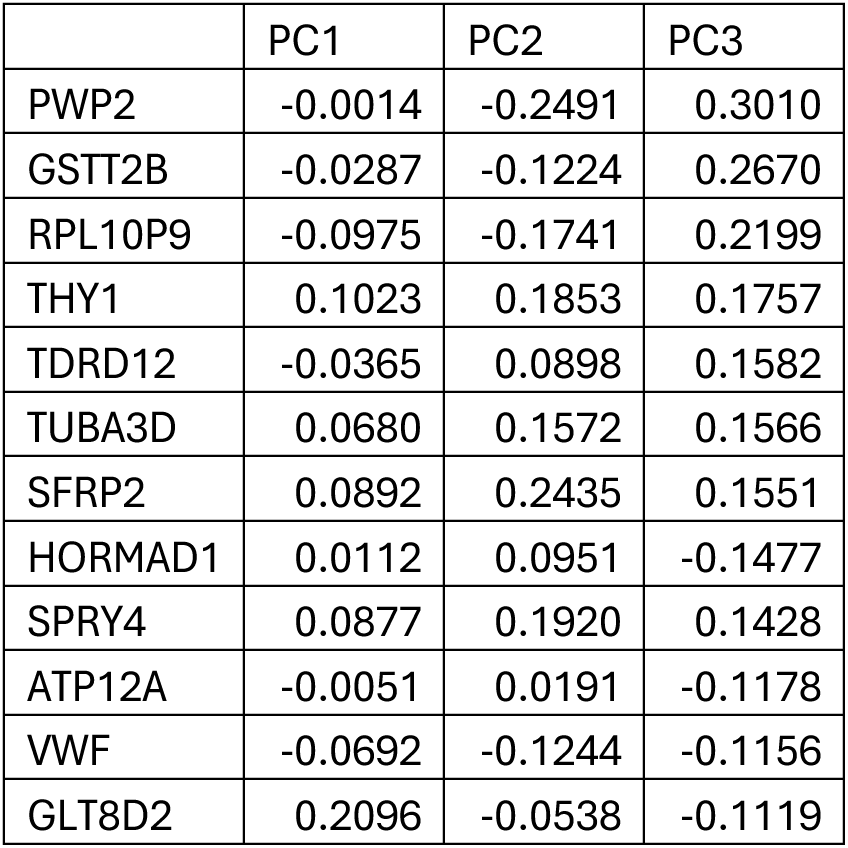
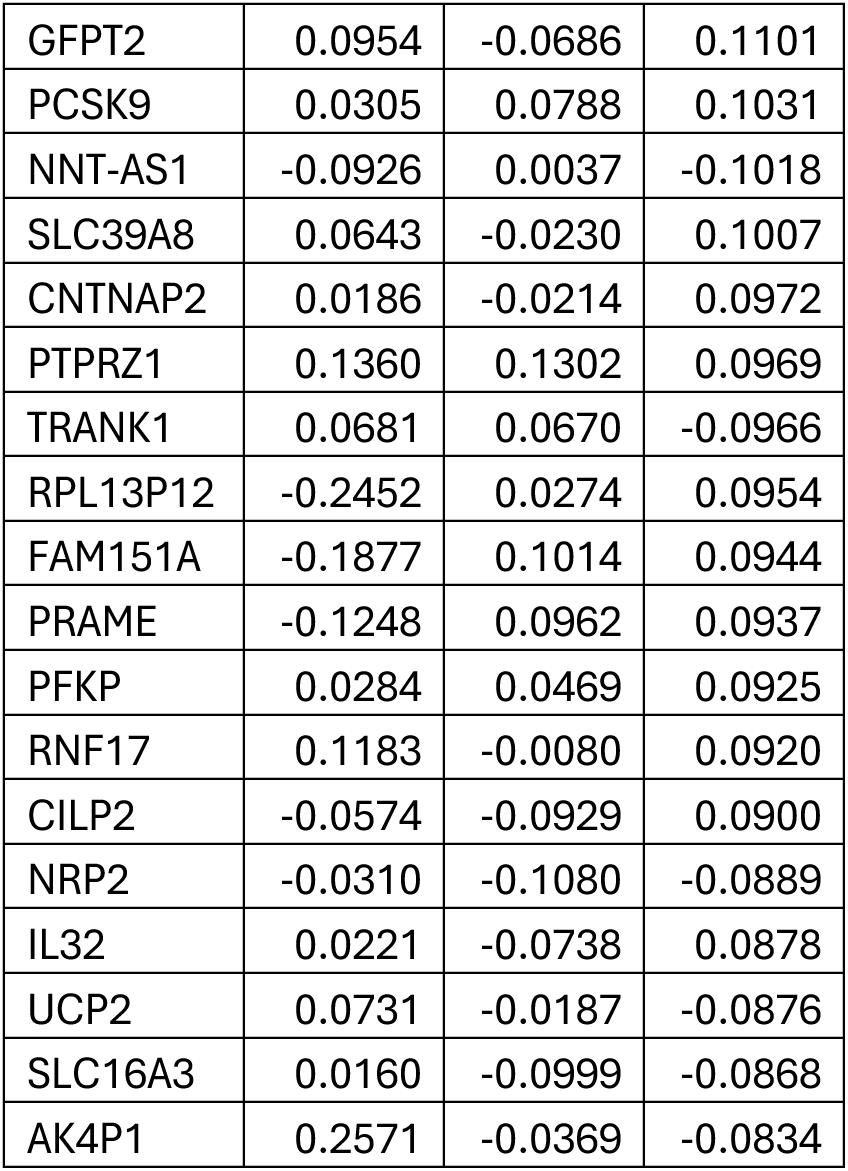
Genes contributing to separation between t2iLGöXYaa and PGXL by PCA Contribution to the first three principal components from PCA analysis of cells cultured in t2iLGöXYaa or PGXL. 30 genes with the greatest absolute contribution to PC3 are included.

|  | PC1 | PC2 | PC3 |
| --- | --- | --- | --- |
| PWP2 | -0.0014 | -0.2491 | 0.3010 |
| GSTT2B | -0.0287 | -0.1224 | 0.2670 |
| RPL10P9 | -0.0975 | -0.1741 | 0.2199 |
| THY1 | 0.1023 | 0.1853 | 0.1757 |
| TDRD12 | -0.0365 | 0.0898 | 0.1582 |
| TUBA3D | 0.0680 | 0.1572 | 0.1566 |
| SFRP2 | 0.0892 | 0.2435 | 0.1551 |
| HORMAD1 | 0.0112 | 0.0951 | -0.1477 |
| SPRY4 | 0.0877 | 0.1920 | 0.1428 |
| ATP12A | -0.0051 | 0.0191 | -0.1178 |
| VWF | -0.0692 | -0.1244 | -0.1156 |
| GLT8D2 | 0.2096 | -0.0538 | -0.1119 |
| GFPT2 | 0.0954 | -0.0686 | 0.1101 |
| PCSK9 | 0.0305 | 0.0788 | 0.1031 |
| NNT-AS1 | -0.0926 | 0.0037 | -0.1018 |
| SLC39A8 | 0.0643 | -0.0230 | 0.1007 |
| CNTNAP2 | 0.0186 | -0.0214 | 0.0972 |
| PTPRZ1 | 0.1360 | 0.1302 | 0.0969 |
| TRANK1 | 0.0681 | 0.0670 | -0.0966 |
| RPL13P12 | -0.2452 | 0.0274 | 0.0954 |
| FAM151A | -0.1877 | 0.1014 | 0.0944 |
| PRAME | -0.1248 | 0.0962 | 0.0937 |
| PFKP | 0.0284 | 0.0469 | 0.0925 |
| RNF17 | 0.1183 | -0.0080 | 0.0920 |
| CILP2 | -0.0574 | -0.0929 | 0.0900 |
| NRP2 | -0.0310 | -0.1080 | -0.0889 |
| IL32 | 0.0221 | -0.0738 | 0.0878 |
| UCP2 | 0.0731 | -0.0187 | -0.0876 |
| SLC16A3 | 0.0160 | -0.0999 | -0.0868 |
| AK4P1 | 0.2571 | -0.0369 | -0.0834 |

**Table S3.**
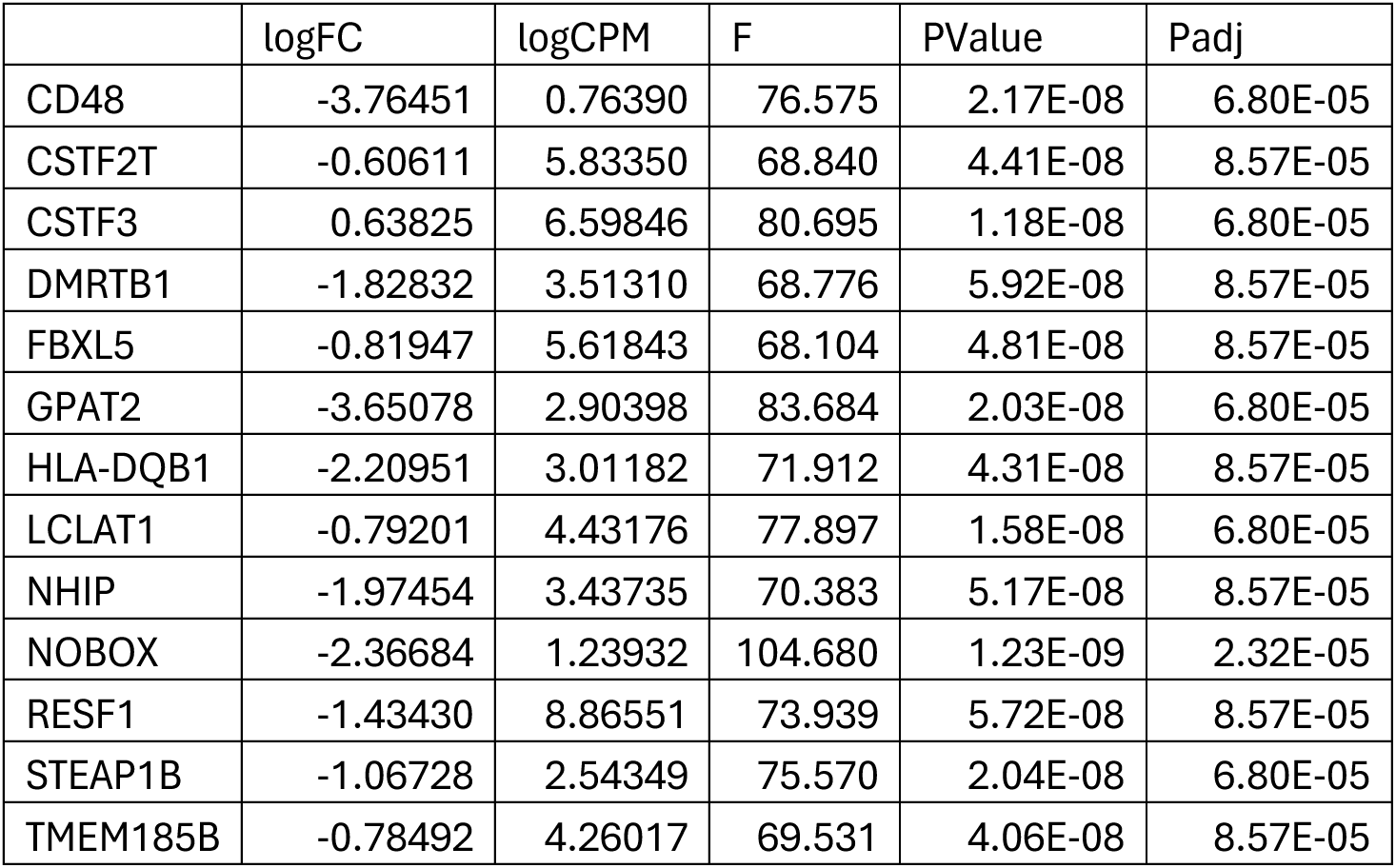
Differentially expressed genes between cells cultured in t2iLGöXYaa and PGXL Log2 fold change (positive values higher in t2iLGöXYaa) in expression and log2 mean expression for factors the 13 genes found to be differentially expressed (Padj < 0.0001) between t2iLGöXYaa and PGXL culture.

**Table S4.**
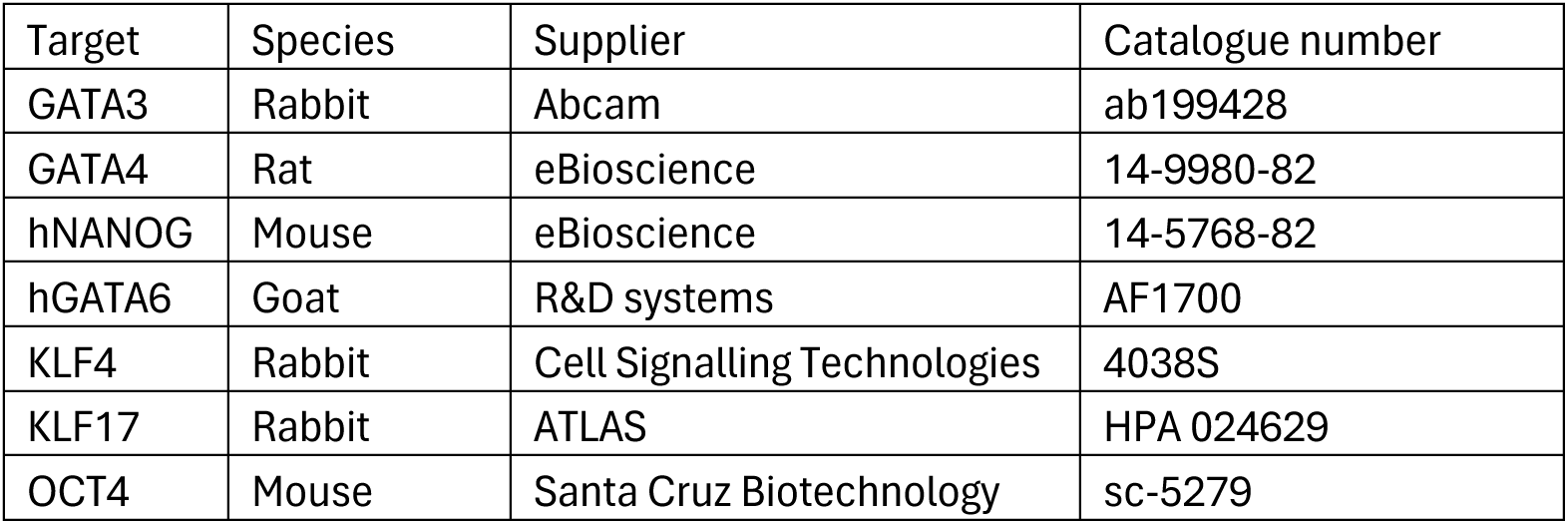
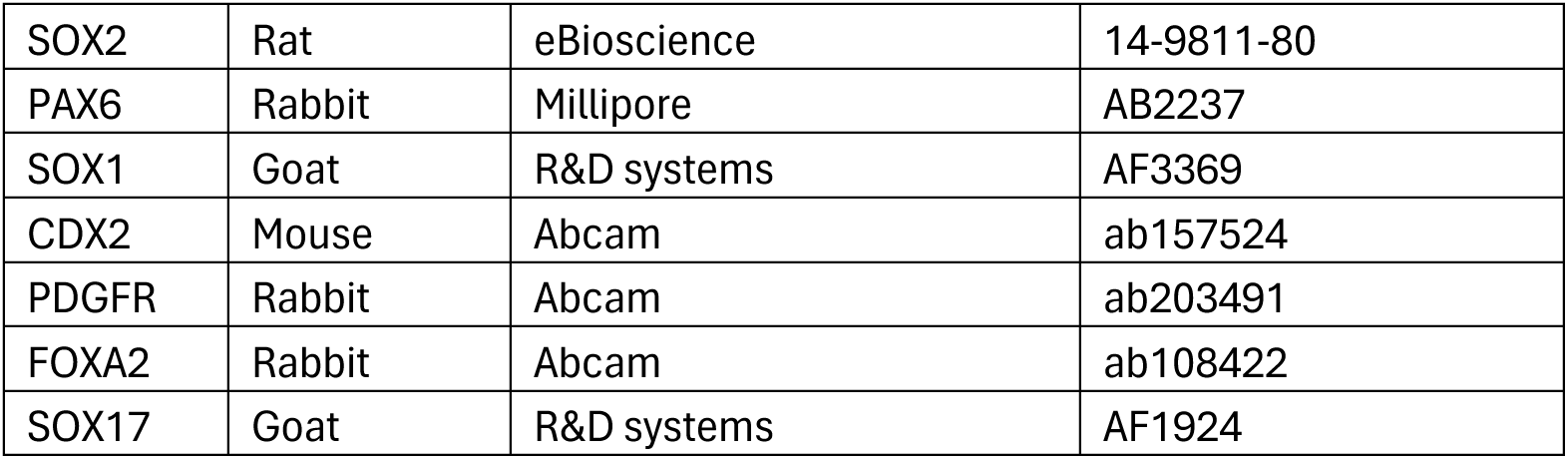
List of primary antibodies used in this study.

**Table S5.**
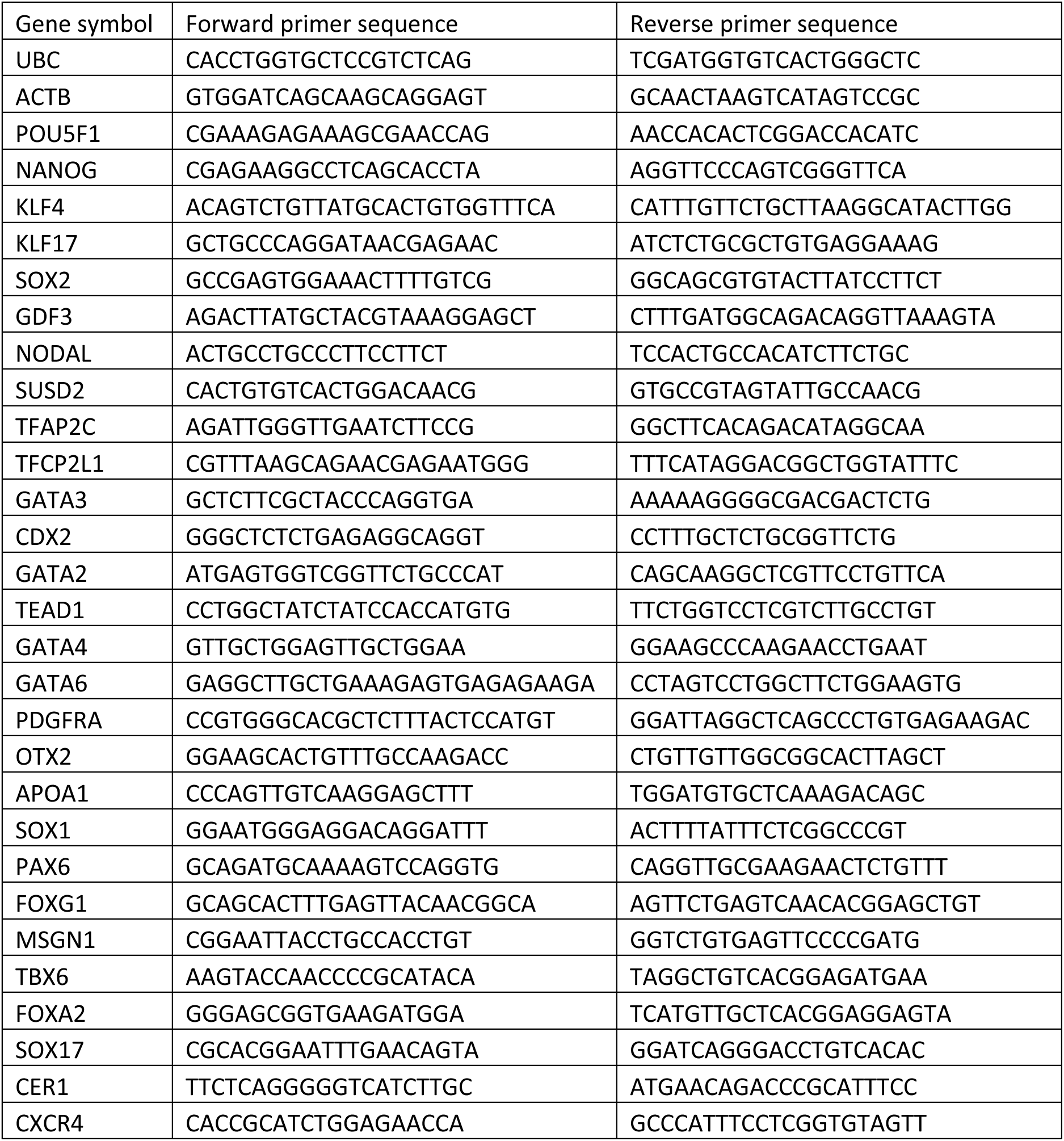
List of RT-qPCR primer sequences used in this study.

